# Vitamin B12 status and folic acid supplementation influence mitochondrial heteroplasmy levels in mice as they age

**DOI:** 10.1101/2023.06.15.545050

**Authors:** Darren J Walsh, David J Bernard, Joanna L Fiddler, Faith Pangilinan, Madison Esposito, Denise Harold, Martha S Field, Anne Parle-McDermott, Lawrence C Brody

**Affiliations:** Gene and Environment Interaction Section, National Human Genome Research Institute, NIH, Bethesda, MD, USA; School of Biotechnology, Dublin City University, Dublin, Ireland; Division of Nutritional Sciences, Cornell University, Ithaca NY 14850; Department of Food, Nutrition, and Packaging Sciences, Clemson University, Clemson SC 29634

**Keywords:** heteroplasmy, mitochondria, folic acid, cobalamin, vitamin B12

## Abstract

One-carbon metabolism is a complex network of metabolic reactions that are essential for cellular function including DNA synthesis. Vitamin B12 and folate are micronutrients that are utilized in this pathway and their deficiency can result in the perturbation of one-carbon metabolism and subsequent perturbations in DNA replication and repair. This effect has been well characterized in nuclear DNA but to date, mitochondrial DNA (mtDNA) has not been investigated extensively. Mitochondrial variants have been associated with several inherited and age-related disease states; therefore, the study of factors that impact heteroplasmy are important for advancing our understanding of the mitochondrial genome’s impact on human health.

Heteroplasmy studies require robust and efficient mitochondrial DNA enrichment to carry out in-depth mtDNA sequencing. Many of the current methods for mtDNA enrichment can introduce biases and false positive results. Here we use a method that overcomes these limitations and have applied it to assess mitochondrial heteroplasmy in mouse models of altered one-carbon metabolism. Vitamin B12 deficiency was found to cause increased levels of mitochondrial DNA heteroplasmy across all tissues that were investigated. Folic acid supplementation also contributed to elevated mitochondrial DNA heteroplasmy across all mouse tissues investigated. Heteroplasmy analysis of human data from the Framingham Heart Study suggested a potential sex-specific effect of folate and vitamin B12 status on mitochondrial heteroplasmy. This is a novel relationship that may have broader consequences for our understanding of one-carbon metabolism, mitochondrial related disease and the influence of nutrients on DNA mutation rates.

**Significance Statement:** Using a sensitive method for mitochondrial heteroplasmy analysis, we show that both vitamin B12 and folic acid can impact mitochondrial DNA mutation. This effect requires further investigation of the potential impact on humans.

## Introduction

Cobalamin (vitamin B12) and folate (vitamin B9) are essential micronutrients that animals must obtain from their diet and play an essential role in one-carbon metabolism. Vitamin B12 deficiency can manifest itself in the development of symptoms including impaired cognition, infertility, depression, myelopathy, ataxia and cardiomyopathy^1,2^. It can also contribute to or exacerbate: methylmalonic aciduria, megaloblastic anemia, age-related macular degeneration, improper bone development, autism and schizophrenia^3–7^. Suboptimal folate status during the periconceptional period of pregnancy has an established link with increased risk of neural tube defects (NTDs)^8^, with a number of studies also reporting low or inadequate maternal B12 status as an additional NTD and omphalocele^9,10^ risk factor. Folate deficiency is also associated with increased risk of congenital heart defects, megaloblastic anemia, inflammatory bowel disease (IBS), Parkinson’s disease and cancer^11–16^. Although vitamin B12 and folate deficiency have been associated with these conditions, the exact molecular underpinnings that drive them are not yet fully known.

Given that NTD risk is associated with suboptimal folate status, folic acid fortification has been implemented in countries across the world in an attempt to reduce this risk among the general population^17^. Although the positive impact of folic acid fortification is widely accepted, there is a concern related to over-supplementation. High folic acid intake has been associated with increased risk of a number of cancer types^18–20^. Extremes in folic acid intake have recently been shown to have surprising, complex and tissue-specific effects on mutagenicity. Supplementation caused increased nuclear DNA mutation frequency in the colon, and deficiency caused the same effect in bone marrow in mice^21^. A well-established concern with folic acid supplementation is the masking of vitamin B12 deficiency through the reversal of anemia but with the continued risk of causing irreversible neurological effects due to disrupted cobalamin-dependent reactions^22^. The existence and extent of this effect is subject to debate among experts^22^. Folic acid supplementation has also been associated with increased neurological issues in individuals with vitamin B12 deficiency^23^. An expert panel organized by the US National Toxicology program agreed that there is some limited but supporting evidence of exacerbating neurological problems associated with high folate/folic acid in the presence of vitamin B12 deficiency^24^. This panel also recognized that there is some evidence of folic acid supplementation causing accelerated colon cancer progression in rodents^25^. Although there is some supportive data suggesting that there are potential risks associated with folic acid supplementation, there are many difficulties comparing studies due to variability in the methods used to measure folic acid/folate status.

One molecular mechanism that is potentially driving these micronutrient effects could be through an impact on DNA synthesis and possibly methylation patterns. Vitamin B12 deficiency can inhibit *de novo* DNA synthesis and DNA methylation, which is thought to be a contributing factor to some vitamin B12 deficient phenotypes^26–28^. Global DNA hypomethylation has been reported in mice fed vitamin B12 deficient diets^27^. B12 deficiency also impairs *de novo* thymidylate synthesis, which leads to uracil misincorporation into DNA^29–33^. There are multiple mechanisms through which folate deficiency can impact DNA, such as damage due to nucleotide misincorporation including uracil, an increase in reactive oxidative species^34–36^ and epigenetic effects that may result in methylation changes^37^. However, many of the studies that have investigated the effect of vitamin B12 and folate status on DNA integrity have primarily focused on nuclear DNA, which does not consider the mitochondrial genome and the unique impact that disruption to mtDNA synthesis may have on cellular function.

Unlike the diploid nuclear genome, the fluctuating, multi-copy nature of mtDNA means that nucleotide variants can be present at any frequency within a cell. The presence and frequency of mtDNA variants is referred to as mitochondrial heteroplasmy. Elevated heteroplasmy at particular locations within the mitochondrial genome has been linked to several age-related and mitochondrial-related diseases such as Leber hereditary optic neuropathy, mitochondrial myopathies, Parkinson’s disease, coronary artery disease and hypertension^38–45^. The exact relationship between heteroplasmy and the manifestation of these biological features is not yet known and is further complicated by complex penetrance, tissue-specific heteroplasmy and a lack of understanding of how particular single nucleotide variants may impact mitochondrial functionality. The pathogenic heteroplasmic variants linked to these conditions are routinely found in unaffected individuals, however the frequency of said variants is usually lower in these circumstances^46^. The aim of this paper was to investigate the impact of vitamin B12 and folate status on mtDNA heteroplasmy. We find that both vitamin B12 status and folic acid supplementation has an impact on mtDNA mutation rates across all investigated tissues in rodent models. Further, targeted studies are needed to assess this effect in humans.

## Results

### Investigation of age and vitamin B12 status on mtDNA heteroplasmy in transcobalamin receptor mutant mice using Mito-SiPE

In mice and humans, vitamin B12 is distributed to tissues through the blood in a complex with transcobalamin (TC), a secreted protein encoded by transcobalamin 2 gene^47^. Transcobalamin transports B12 into celsl by interacting with its receptor which is encoded by *Cd320*. Our lab previously developed and characterized a viable *Cd320*^−/−^ mouse model that mimics vitamin B12 deficiency in humans^48^. These mice presented hallmarks consistent with that of human vitamin B12 deficiency: increased plasma methylmalonic acid (MMA) and plasma homocysteine (Hcy) concentrations. The phenotype of these mice can be further exacerbated via dietary vitamin B12 restriction which produces infertility in female mice and fatal macrocytic anemia in mice after approximately 1 year on a vitamin B12 deficient diet^48^. These genetic and dietary manipulations are designed to perturb one carbon metabolism and serve as a model to study vitamin B12 deficiency. Using our previously described methodology, Mito-SiPE^49^, we sequenced the mitochondrial DNA of the aforementioned *Cd320*^−/−^ mice to ask whether our genetic and/or dietary manipulations influenced heteroplasmy. We analyzed seven different tissues: brain, heart, lungs, spleen, liver, kidney and muscle in mice maintained for approximately one year on vitamin B12 deficient or replete diets.

#### Mitochondrial heteroplasmy was elevated in older mouse tissues compared to younger tissues

We assessed heteroplasmy using three interrelated metrics: 1) the number of heteroplasmic sites, 2) the average alternative allele frequency (average heteroplasmy) and 3) the cumulative heteroplasmic burden. The number of heteroplasmic sites is the number of nucleotide positions at which an alternative allele was identified above the threshold frequency (0.2%) in a sample. Average heteroplasmy is the mean frequency of all variants observed in a sample. Finally, cumulative heteroplasmic burden is the sum of all variant frequencies that were identified above the threshold frequency in a sample. These metrics of mitochondrial heteroplasmy have been utilized in previous studies^42,49–54^.

Using these metrics, we observed a significant increase in mitochondrial heteroplasmy in older mice (24 months old) compared with younger mice (12 months) (Fig. 1). The tissues of older mice had a significant increase in the number of heteroplasmic sites (1.59 ± 0.73 sites, mean ± SD) compared to younger mouse tissues (0.27 ± 0.53 sites, mean ± SD, Fig. 1, a). Elevated average heteroplasmy levels were also found in older mice (0.04 ± 0.03, mean ± SD) in contrast to younger mice (0.001 ± 0.001, mean ± SD, Fig. 1, b). Finally, cumulative heteroplasmic burden showed a similar pattern to the previous two metrics with elevated levels in older tissues compared to younger (0.068 ± 0.073, 0.001 ± 0.001, respectively. Fig. 1, c) There was no evidence of tissue-specific trends in heteroplasmy across any of the three metrics (Supplementary Fig. 1).

**Figure 1.**
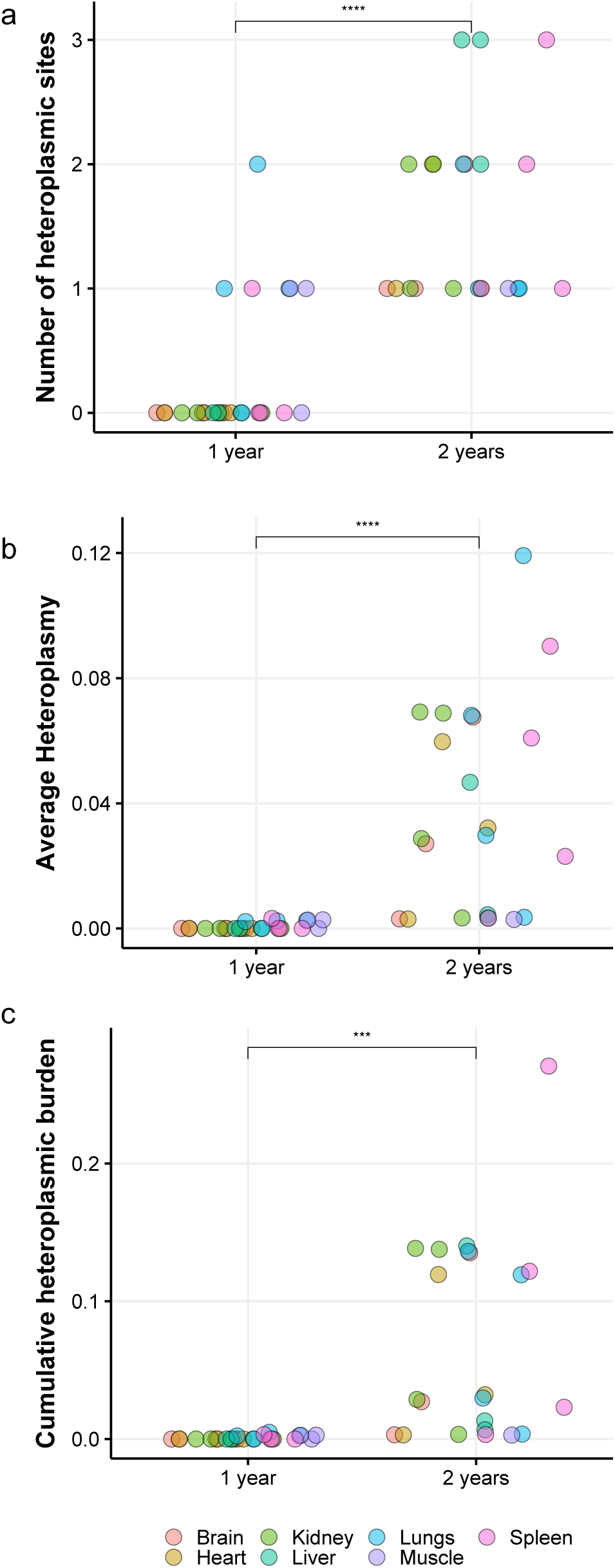
Mitochondrial heteroplasmy is elevated in older mouse tissues. The number of heteroplasmic sites is the amount of nucleotide positions at which an alternative allele was observed in a given sample. a, only 7 unique heteroplasmic variants were found across all tissues of the young cohort. b, Average heteroplasmy is the average frequency of alternative alleles that were observed in a sample. c, Finally, cumulative heteroplasmic burden is the additive frequency of all variants found in one sample. Across the three aforementioned metrics, there was a significant increase in the older mouse tissues (Student’s T-test, P values shown above each plot. *** = P < 0.0005, **** = P < 0.00005). The colors (see key) represent the tissue from which each mtDNA sample was taken from. The number of samples were as follows; 1 year (n=26) and 2 years (n=22); brain (n=4 and n=3, respectively), heart (n=4 and n=3, respectively), kidney (n=4 and n=4, respectively), liver (n=3 and n=3, respectively), lungs (n=4 and n=4, respectively), muscle (n=3 and n=1, respectively) and spleen (n=4 and n=4, respectively).

#### Vitamin B12 deficiency leads to increased mtDNA heteroplasmy

Higher levels of mitochondrial heteroplasmy were observed in mice fed the vitamin B12 deficient diet, regardless of their *Cd320* genotype (two-way ANOVA, Table 1). A significant relationship was observed between diet and number of heteroplasmic sites, average heteroplasmy and cumulative heteroplasmic burden. Both *Cd320*^−/−^ and *Cd320*^+/+^ mice that were fed vitamin B12 deficient chow had statistically significant increases in heteroplasmic sites, average heteroplasmy and cumulative heteroplasmic burden in their tissues compared to controls (Fig. 2). Genotype appeared to have an impact on the number of heteroplasmic sites but not cumulative heteroplasmic burden or average alternative allele frequency. There was no evidence of a tissue specific effect of heteroplasmy (Supplementary Fig. 2).

**Figure 2.**
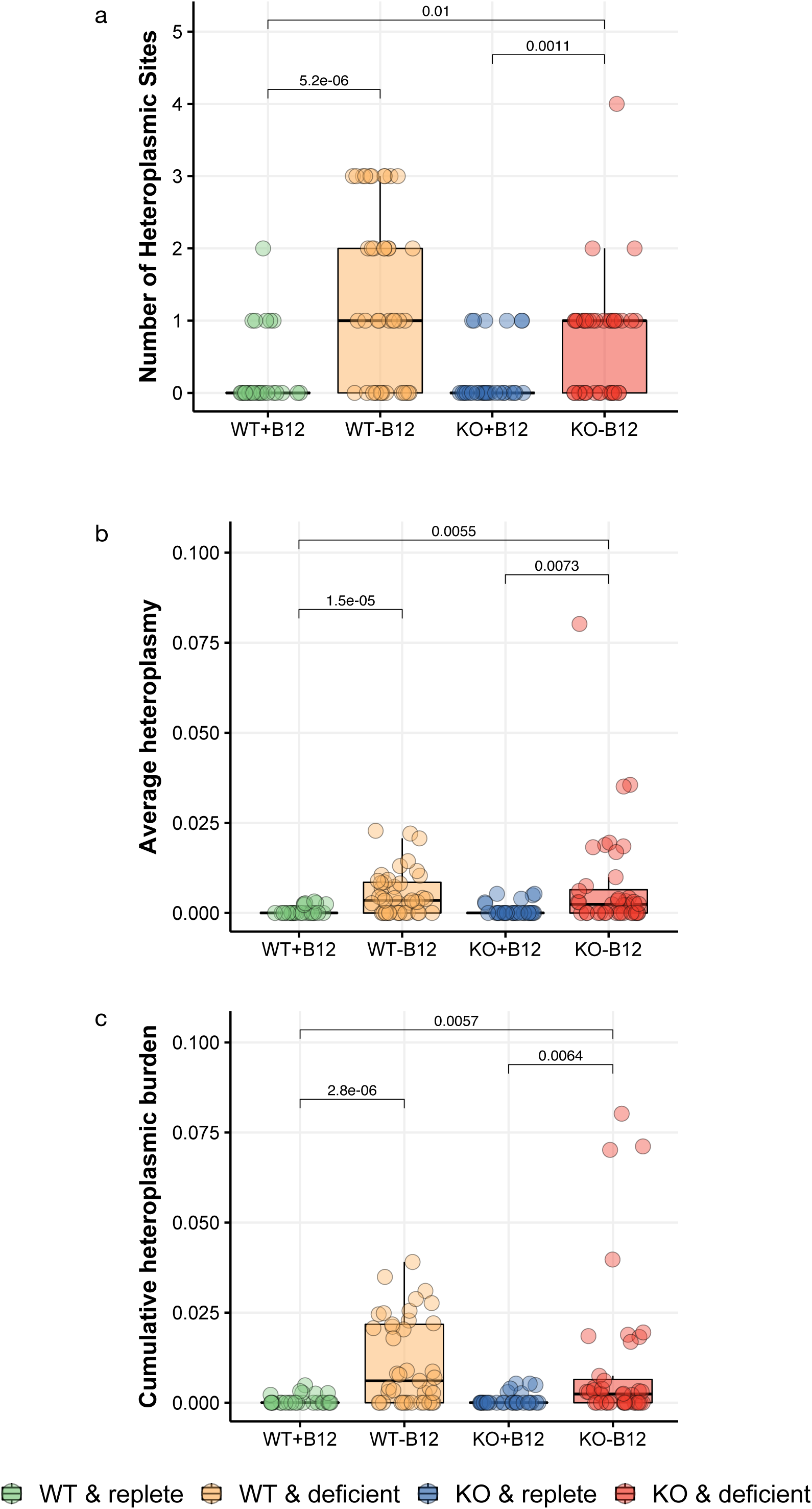
Mitochondrial heteroplasmy was higher in the tissues of mice that were fed a vitamin B12-deficient diet, irrespective of genotype. Elevated levels of heteroplasmy were observed in both Cd320^−/−^ and Cd320^+/+^ mice that were fed the vitamin B12-deficient diet. There was a significant increase in a, the number of heteroplasmic sites, b, average heteroplasmy and c, cumulative heteroplasmic burden (Tukey’s HSD, P values presented above each comparison. Corrected for multiple comparisons using Bonferroni correction). The number of samples were as follows; WT+vitamin B12 (n=26, brain=4, heart=4, kidney=4, liver=3, lungs=4, muscle=3 and spleen=4), WT-vitamin B12 (n=41, brain=6, heart=6, kidney=6, liver=5, lungs=7, muscle=4 and spleen=7), KO+vitamin B12 (n=30, brain=5, heart=5, kidney=5, liver=4, lungs=5, muscle=1 and spleen=5) and KO-vitamin B12 (n=40, brain=7, heart=7, kidney=5, liver=5, lungs=6, muscle=3 and spleen=7).

**Table 1.**
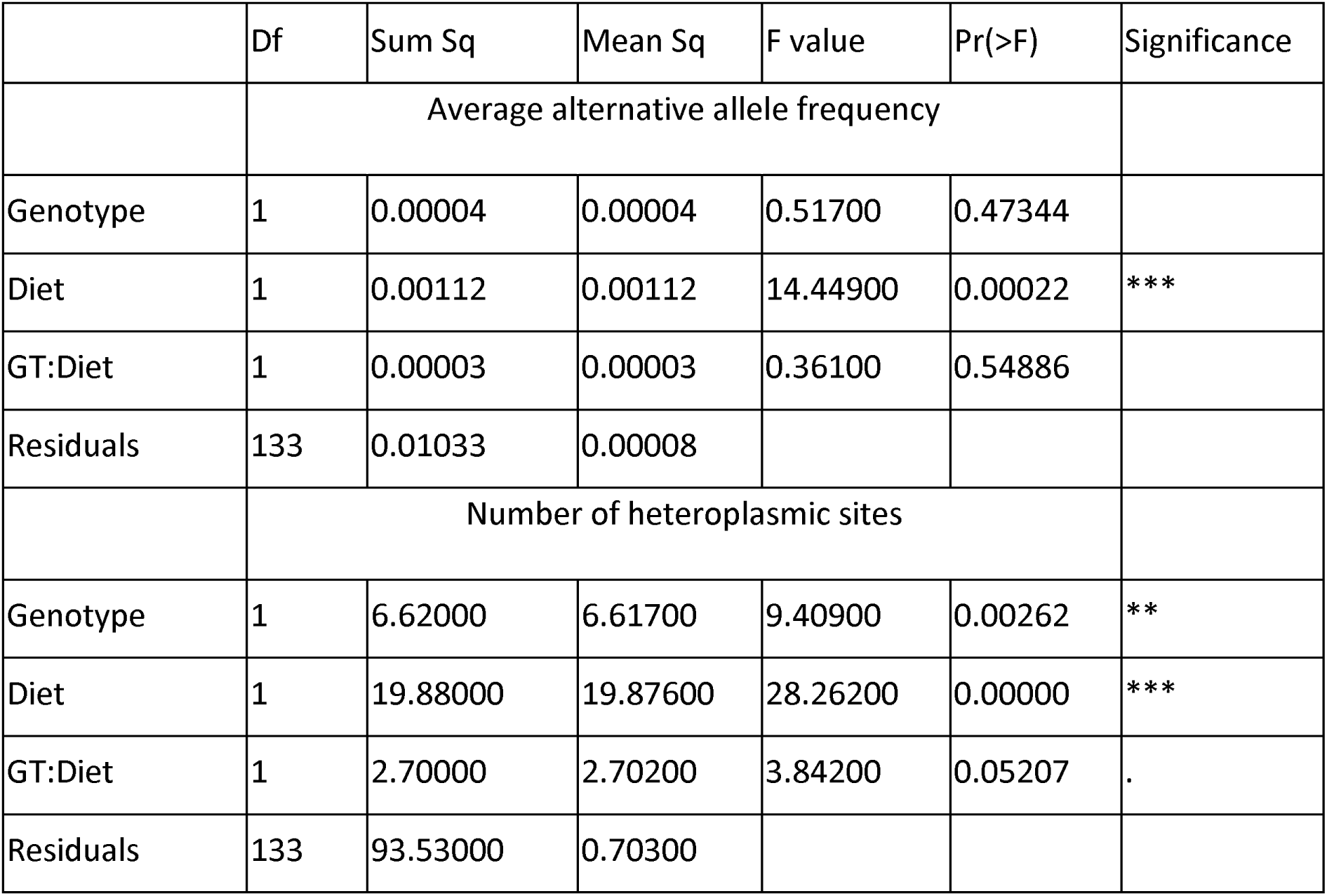
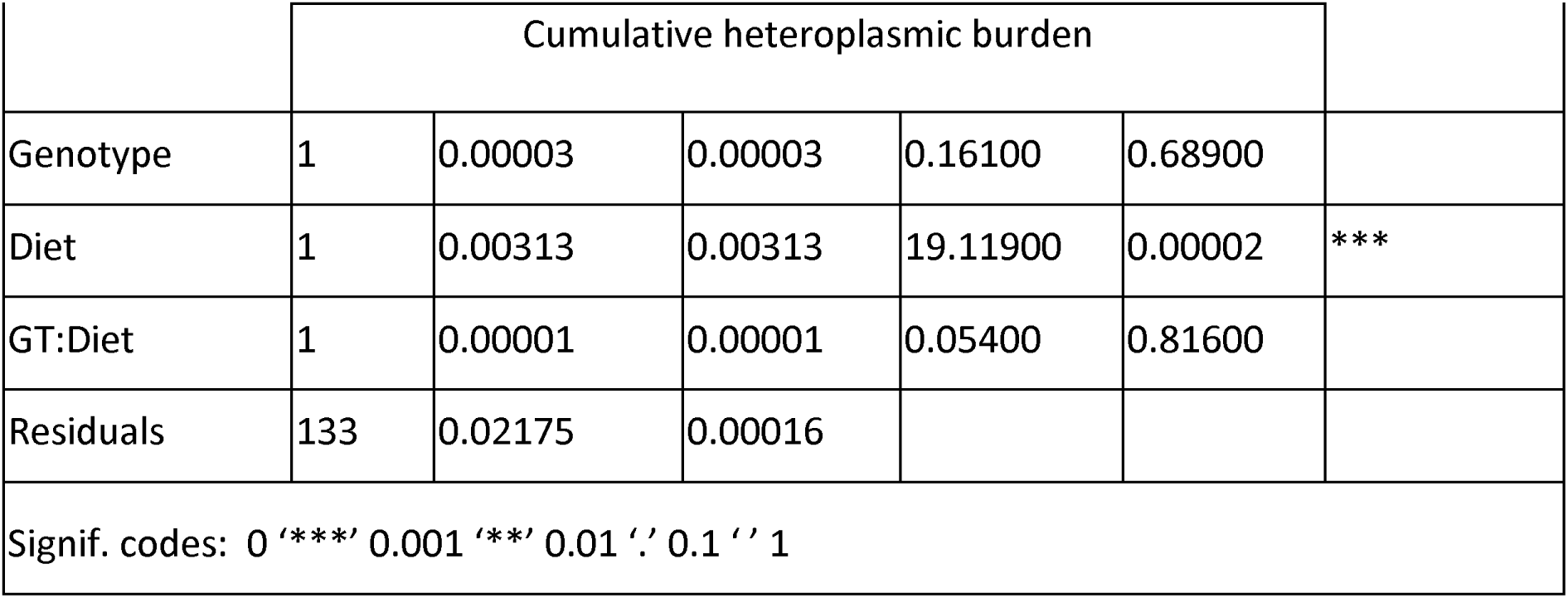
Two-way ANOVA results from analyzing the effects of Cd320 genotype and a vitamin B12 replete or deficient diet on measures of mitochondrial heteroplasmy in mice.

|  | Df | Sum Sq | Mean Sq | F value | Pr(>F) | Significance |
| --- | --- | --- | --- | --- | --- | --- |
|  | Average alternative allele frequency |  |  |  |  |  |
| Genotype | 1 | 0.00004 | 0.00004 | 0.51700 | 0.47344 |  |
| Diet | 1 | 0.00112 | 0.00112 | 14.44900 | 0.00022 | *** |
| GT:Diet | 1 | 0.00003 | 0.00003 | 0.36100 | 0.54886 |  |
| Residuals | 133 | 0.01033 | 0.00008 |  |  |  |
|  | Number of heteroplasmic sites |  |  |  |  |  |
| Genotype | 1 | 6.62000 | 6.61700 | 9.40900 | 0.00262 | ** |
| Diet | 1 | 19.88000 | 19.87600 | 28.26200 | 0.00000 | *** |
| GT:Diet | 1 | 2.70000 | 2.70200 | 3.84200 | 0.05207 | . |
| Residuals | 133 | 93.53000 | 0.70300 |  |  |  |

|  | Cumulative heteroplasmic burden |  |  |  |  |  |
| --- | --- | --- | --- | --- | --- | --- |
| Genotype | 1 | 0.00003 | 0.00003 | 0.16100 | 0.68900 |  |
| Diet | 1 | 0.00313 | 0.00313 | 19.11900 | 0.00002 | *** |
| GT:Diet | 1 | 0.00001 | 0.00001 | 0.05400 | 0.81600 |  |
| Residuals | 133 | 0.02175 | 0.00016 |  |  |  |
| Signif. codes: 0 '***' 0.001 '**' 0.01 '.' 0.1 ' ' 1 |  |  |  |  |  |  |

#### Mitochondrial DNA uracil content assessment

Uracil misincorporation was assessed in the mitochondrial DNA isolated from the liver of mice used in the experiment above. There was no difference in the relative level of uracil content in liver mtDNA of young and old mice (Fig. 3, a). There was a significant relationship between uracil content and experimental group at 1 year old across the whole mitochondrial genome (Table 2, two-way ANOVA. Fig. 3, b paired T-test).

**Figure 3.**
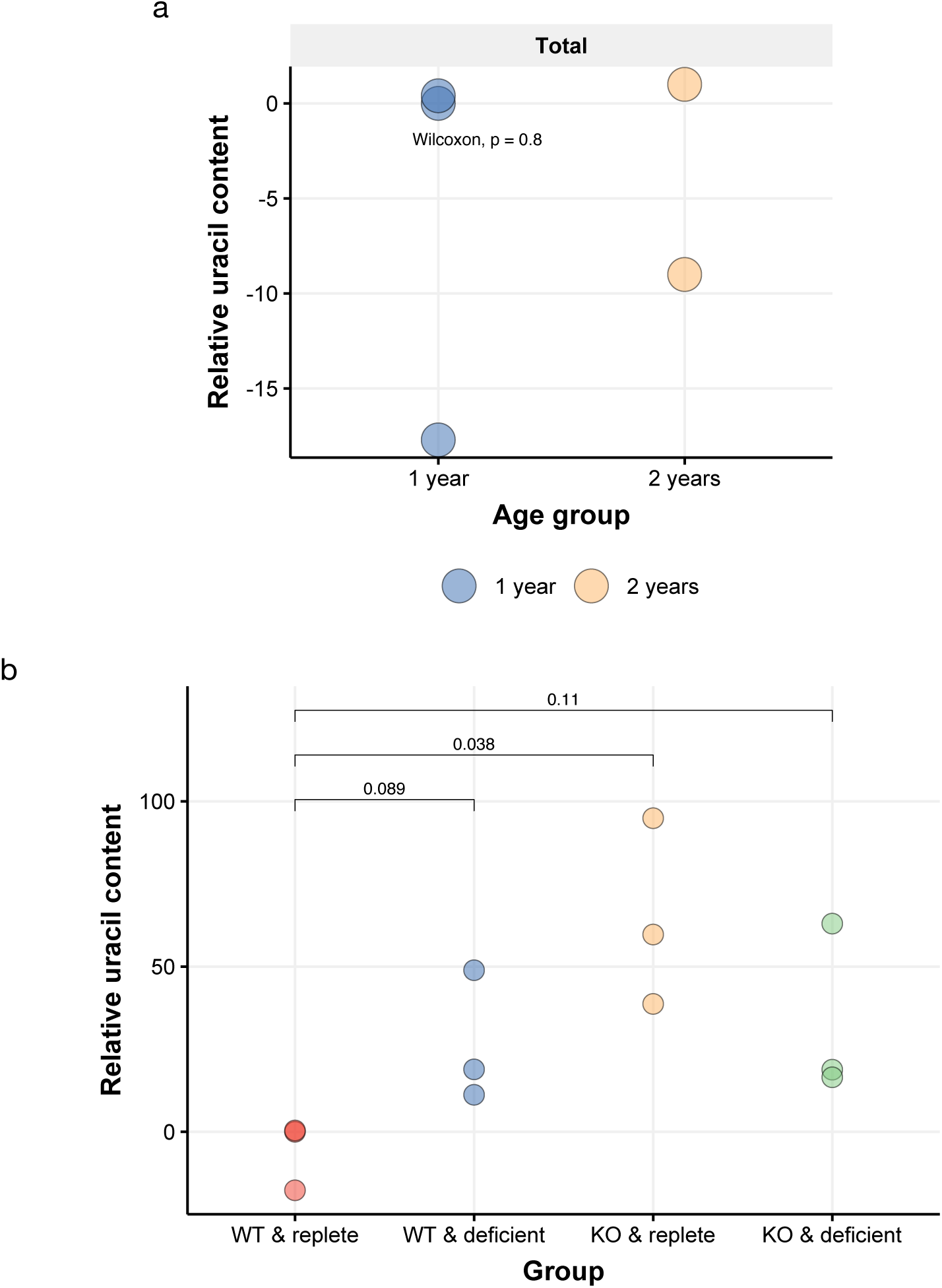
Relative uracil content of liver mitochondrial DNA. No difference in uracil content was observed in liver DNA of young and old mice across any of the six regions or when all regions were combined (a, Wilcoxon rank-sum test). When regions were combined there was a significant relationship between uracil content and experimental group (b, paired Student’s T-test). There were three samples in each experimental group. The y axis represents the relative uracil content. This value was calculated by normalizing a random control sample to zero.

**Table 2.**
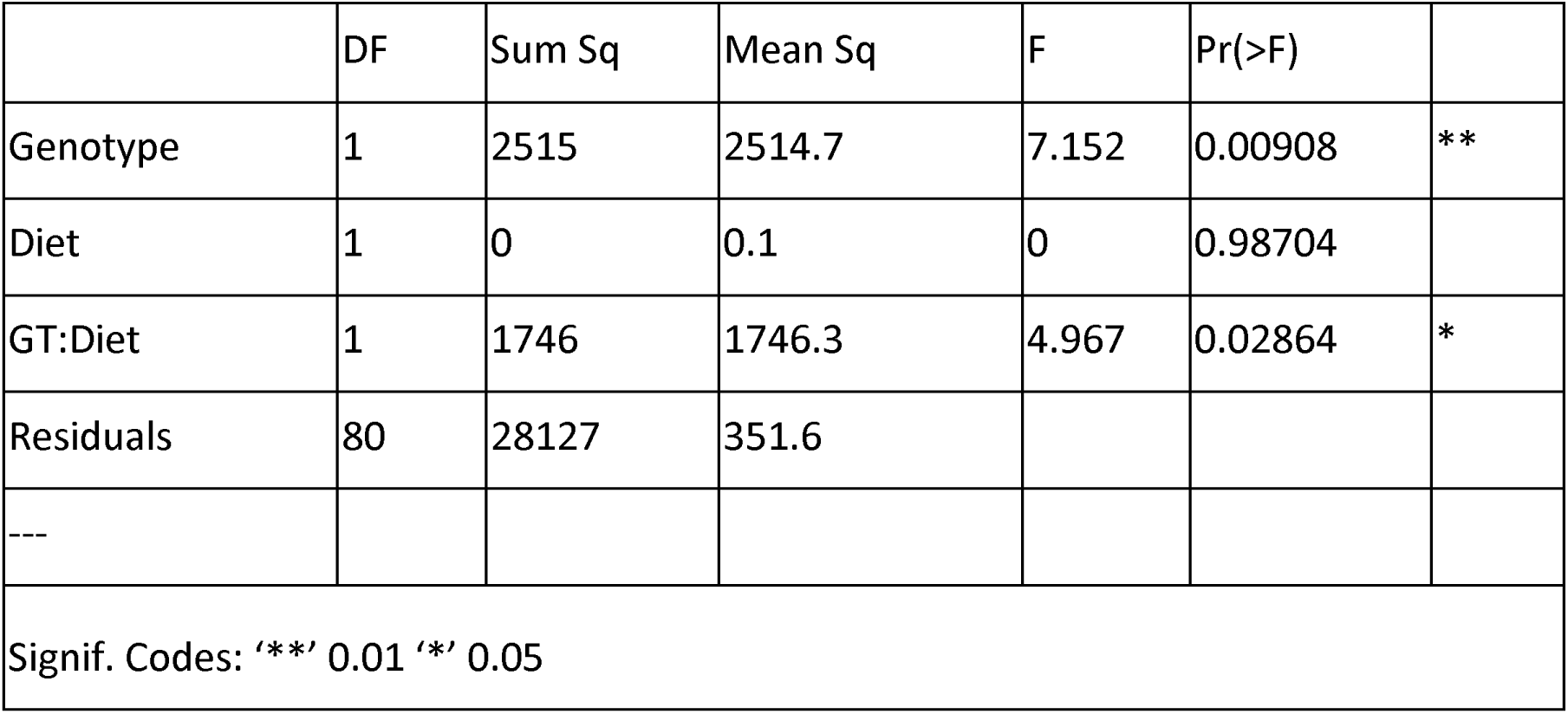
Two-way ANOVA results to measure the effect of folic acid supplementation and vitamin B12 status on measures of mitochondrial heteroplasmy in mice.

### The impact of vitamin B12 status and/or folic acid supplementation on mtDNA heteroplasmy in wild type mice

Tissues of mice that were fed a high folic acid or a vitamin B12 deficient diet appeared to have elevated levels of mtDNA heteroplasmy, in terms of number of heteroplasmic variants (Fig. 4, a), average alternative allele frequency (Fig. 4, b) and cumulative heteroplasmic burden (Fig. 4, c) when compared to the control group (two-way ANOVA, Table 3). Folic acid supplementation had an impact on average alternative allele frequency, number of heteroplasmic sites and cumulative heteroplasmic burden, regardless of vitamin B12 status (Table 3). Vitamin B12 deficiency affected the number of heteroplasmic sites and there appeared to be an effect of the interaction between vitamin B12 status and folic acid supplementation on the number of heteroplasmic sites and average alternative allele frequency (Table 3). The number of heteroplasmic sites had a small range of values across all samples (0-4). Mouse tissues from the control group had 0.15 ± 0.65 heteroplasmic sites and a mean alternative allele frequency of 0.0005 ± 0.003. All the experimental groups had higher levels of heteroplasmy than mice fed a vitamin B12 replete diet with normal levels of folic acid. Interestingly, folic acid supplementation, regardless of vitamin B12 status, appeared to cause an increase in heteroplasmy as did the vitamin B12 deficient diet by itself. There was no obvious tissue-specific effect in average heteroplasmy between the dietary groups (Supplementary Fig. 3). All mice used for the 6-month time-point were male. Both female and male mice were used for heteroplasmy analysis after 12 months on diet (Supplementary Fig. 4). Congruent with the previous results, both sexes in vitamin B12 deficient and folic acid supplemented diets showed elevated levels of mitochondrial heteroplasmy.

**Figure 4.**
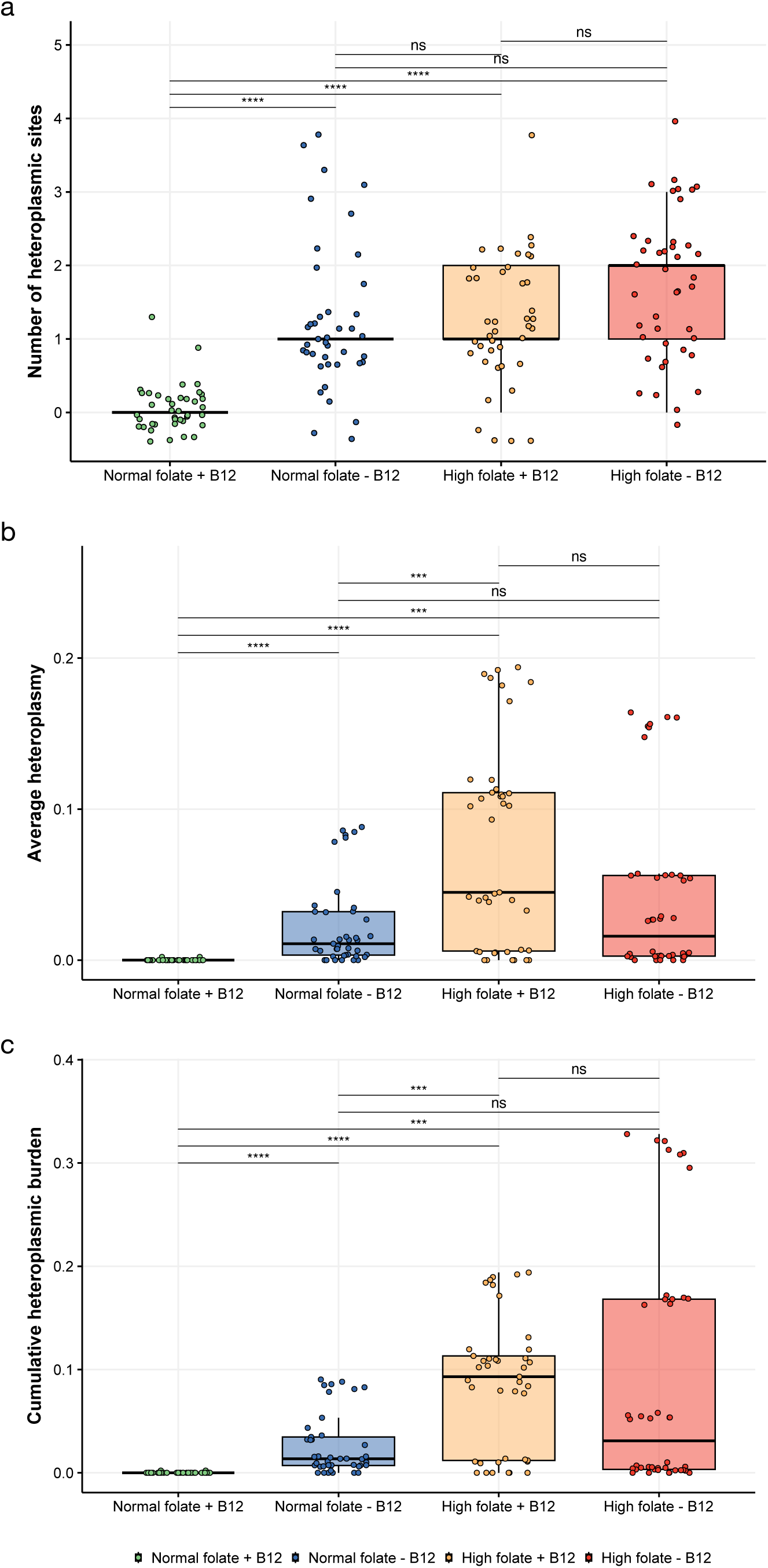
Elevated heteroplasmy levels were observed in the tissues of mice with altered folate/vitamin B12 diet. The average alternative allele frequency was plotted in boxplot format with an overlaid scatterplot. The y axis represents a, average heteroplasmy or alternative allele frequency, b, number of heteroplasmic sites and c, cumulative heteroplasmic burden. The different experimental groups are plotted on the x axis and each point represents one tissue from one mouse. Student’s T-test, Bonferroni corrected P values shown above each plot. *** = P < 0.0005, **** = P < 0.00005.

**Table 3.** Generalized linear models were used to measure the effect size of folate, vitamin B12, age and sex on mitochondrial heteroplasmy in Groups 1 and 2. Gaussian models were used to assess the relationship between folate/vitamin B12 status on average minor allele frequency.

|  | Df | Sum Sq | Mean Sq | F value | Pr(>F) | Significance |
| --- | --- | --- | --- | --- | --- | --- |
|  | Average alternative allele frequency |  |  |  |  |  |
| Folic acid | 1 | 0.0895 | 0.08949 | 42.357 | 9.32E-10 | *** |
| B12 | 1 | 0.0013 | 0.0013 | 0.617 | 0.433147 |  |
| FA:B12 | 1 | 0.0315 | 0.03149 | 14.903 | 0.000164 | *** |
| Residuals | 160 | 0.3381 | 0.00211 |  |  |  |
|  | Number of heteroplasmic sites |  |  |  |  |  |
| Folic acid | 1 | 26.43 | 26.428 | 39.37 | 3.15E-09 | *** |
| B12 | 1 | 28.07 | 28.066 | 41.81 | 1.16E-09 | *** |
| FA:B12 | 1 | 6.9 | 6.898 | 10.28 | 0.00163 | ** |
| Residuals | 160 | 107.41 | 0.671 |  |  |  |
|  | Cumulative heteroplasmic burden |  |  |  |  |  |
| Folic acid | 1 | 0.2294 | 0.22944 | 48.435 | 8.31E-11 | *** |
| B12 | 1 | 0.0103 | 0.01026 | 2.166 | 0.143 |  |
| FA:B12 | 1 | 0.0043 | 0.00433 | 0.914 | 0.341 |  |
| Residuals | 160 | 0.7579 | 0.00474 |  |  |  |

### The impact of vitamin B12 and folate status on mtDNA heteroplasmy in a human cohort

To determine whether B12 and folate status might impact mtDNA heteroplasmy in humans, data from the Framingham Heart Study(FHS)^55^ were analyzed. Briefly, the FHS is a large, longitudinal study in the USA, that began in 1948 with the initial goal of studying cardiovascular disease. The FHS study has evolved over the study’s duration allowing us to use some of the data in our current study investigating the effect of one-carbon metabolism on mtDNA heteroplasmy in humans. There are two sub-cohorts within the larger FHS whose blood folate and vitamin B12 were measured using different methods (hereby referred to as Group 1 and Group 2, respectively. Supplementary Table 3). The methodology used for each of these measurements is described in Materials and Methods. Of these individuals, another smaller subset has whole genome and/or whole exome sequencing (WGS and WXS, respectively) data also available. The age of each participant at their induction to the study was available, although the exact dates for which the blood draws and sequencing were performed were not available for our analyses. This is a limitation of the dataset, as the age at induction will correlate with the age at which tests were performed but it will not be the exact age of each participant at the time of blood draw.

#### Assessment of DNA sequencing data and coverage

The sequencing coverage of the mitochondrial genome was substantially higher in samples that had whole genome sequencing data compared to those with whole exome sequencing. There was no clear difference in read depth across the mitochondrial genome between men and women. In this human data, minor allele frequency was assessed in contrast to alternative allele frequency which was assessed in mice. This is due to the genetic variation that was present in the human data which was not present in the mouse data.

#### Correlation between folate, vitamin B12 and mtDNA heteroplasmy

Folate and vitamin B12 measurements were taken in 1,991 participants in group 1, 157 of which had sequencing data available (Supplementary Table 3). After filtering for a minimum coverage of 100X, 58 participants had heteroplasmic sites identified (Range 1-12). Generalized linear models were used to measure the effect size of folate, vitamin B12, age and sex on mitochondrial heteroplasmy in group 1 (Table 4). Gaussian models were used to assess the relationship between folate/vitamin B12 status on average minor allele frequency. A Poisson regression model was used to assess the number of heteroplasmic sites. Folate concentration was found to have a nominally significant impact on minor allele frequency using a Gaussian model (β= 0.0037, CI= 0.0004 to 0.0070, P=0.0309). Both sex and vitamin B12 concentration influenced the number of heteroplasmic sites (β= −0.3503, CI= −0.6563 to −0.0472 and β= −0.0007, CI= −0.0012 to −0.0001, P=0.0225, respectively. Poisson regression).

**Table 4.**
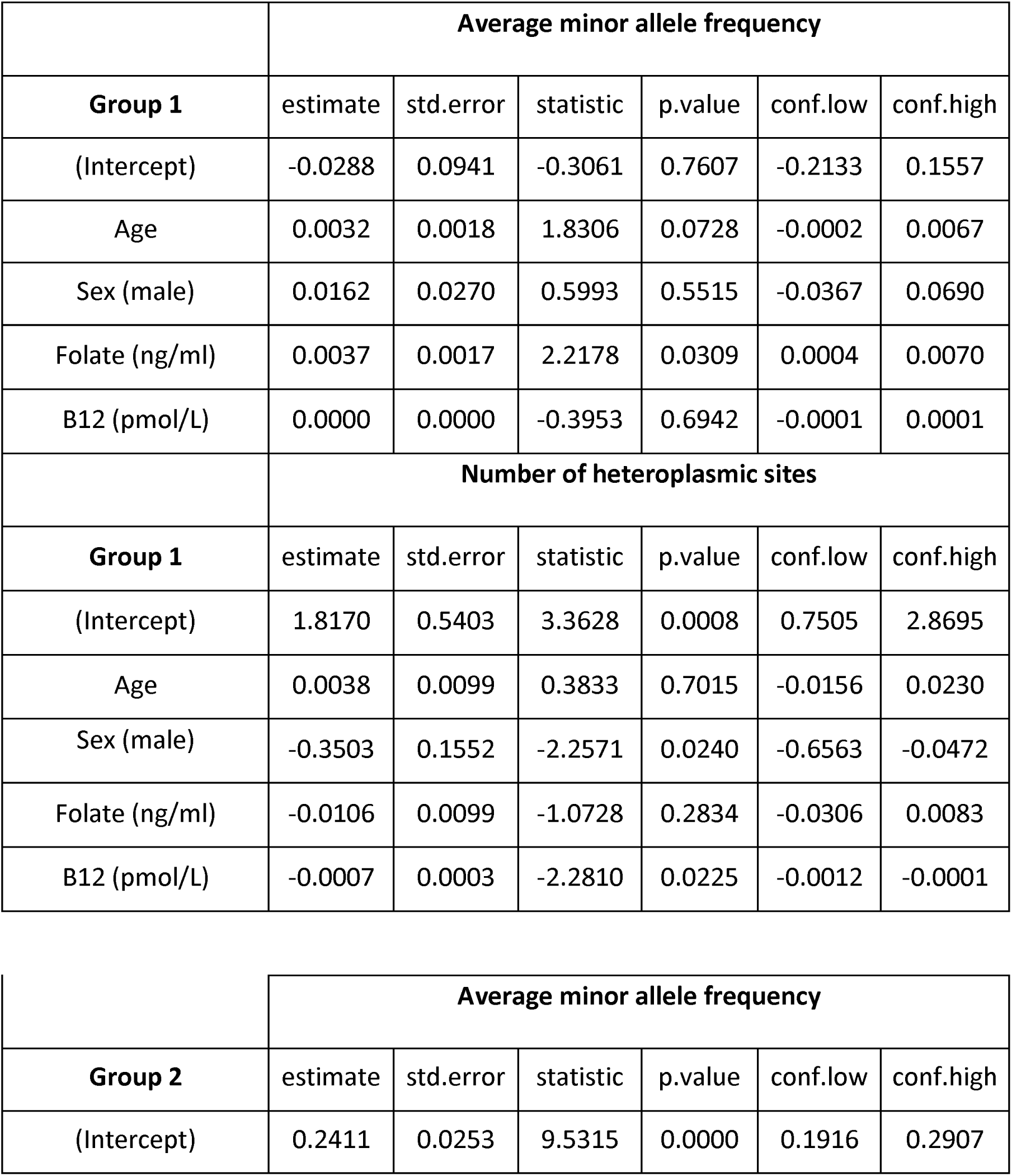
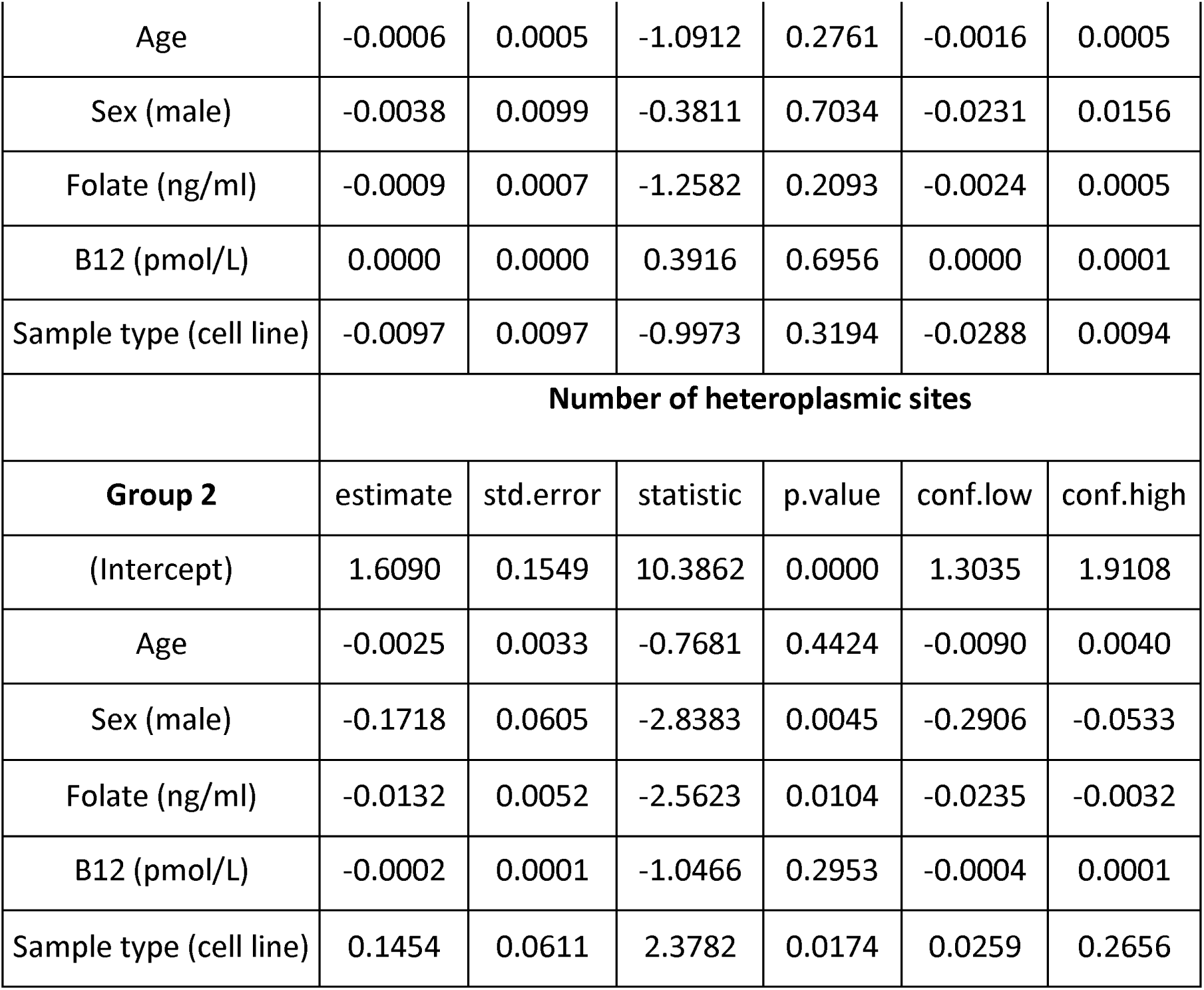
predicted effect-sizes using generalized linear models for average minor allele frequency and number of heteroplasmic sites in Group 1 and Group 2 of the Framingham Heart Study.

The number of individuals in Group 2 with heteroplasmy calls and biochemical measurements post-filtering was 293 (Supplementary Table 3). Whole genome sequencing was performed on DNA originating from blood or cell line samples (n=133 and n=160 post-filtering, respectively. Supplementary Table 3). Age did not correlate with elevated mitochondrial heteroplasmy either in terms of average minor allele frequency or number of heteroplasmic sites in men or women in Group 2 (Table 4). Folate concentration had no impact on average minor allele frequency but did influence the number of heteroplasmic sites (β= −0.0132, CI= −0.0235 to −0.0032, Table 4). Mitochondrial heteroplasmy did not appear to be associated with plasma vitamin B12 concentrations when average minor allele frequency and number of heteroplasmic sites were considered (Table 4). Sample type and sex were both found to have a significant impact on the number of heteroplasmic sites (®=0.1454, CI= 0.0259 to 0.2656, P=0.0174 and ®= −0.1718, CI= −0.2906 to –0.0533, P=0.0045, respectively).

## Discussion

The aim of this study was to determine if conditions that impact one-carbon metabolism had an effect on mtDNA heteroplasmy. Both vitamin B12 and folate are essential micronutrients in one-carbon metabolism. We found that large changes in nutrients involved in one-carbon metabolism can cause increased heteroplasmy in mice. This effect was less consistent in humans using an observational study design.

### Mitochondrial sequencing coverage, mtDNA heteroplasmy analysis and age

The three metrics we used to assess mitochondrial heteroplasmy have been used in several studies^42,49–54^. While they are related measures, each provides a slightly nuanced view on the level of heteroplasmy in a sample. We found elevated levels of mitochondrial heteroplasmy in 24-month-old mice compared to 12-month-old mice using the three aforementioned metrics. This was expected as the relationship between heteroplasmy and aging in mice and humans has been well characterized^38,54,56–62^. It is surprising however, that this effect was observed across all tissues and not in a tissue-specific manner. This finding is in contrast to most heteroplasmy studies to-date, which almost always record a tissue-specific effect on heteroplasmy^42,49–54^.

We hypothesize that the reported tissue differences may be related to nuclear mitochondrial sequences (NUMT) contamination causing false positive heteroplasmy calls in studies using long-range PCR amplification or probe hybridisation to enrich mtDNA. In these studies, it is possible that changes in mtDNA copy number cause NUMT contamination to be enriched at different levels and falsely identified as heteroplasmy. This would mean that changes in apparent ‘heteroplasmy’ as measured by these sequence dependent methods could be a proxy for mtDNA copy number changes, which would explain tissue-specific changes in heteroplasmy observed in these studies. The Mito-SiPE methodology utilized in this research, whilst not totally eliminating the effects of NUMTs, drastically reduces their impact on subsequent heteroplasmy calls^49^. Notably, only 17 variants were found across all 329 samples above a frequency of 10%. This is somewhat unexpected as many studies to date that have employed sequence-dependent methods, in both humans and mice, have frequently documented variants at higher frequencies^42,50,53,54^.

### Effect of vitamin B12 depletion and folic acid supplementation on mtDNA heteroplasmy

Higher levels of heteroplasmy were observed in mice that were maintained on a vitamin B12 deficient diet, regardless of their transcobalamin receptor mutation (*Cd320)* genotype. Like the differences between young and old mice, there were no obvious tissue-specific effects. Previous studies have noted that altered one-carbon metabolism can cause changes to nuclear DNA, although to the best of our knowledge, our report is the first time that vitamin B12 status has been shown to have a direct effect on mitochondrial heteroplasmy^28,34,63,64^. While the magnitude of this effect appears to be modest, it is noteworthy that the level of change we observed are of the same order as observed between young and old mice. The impact of this level of heteroplasmy frequency on mitochondrial function is unknown. These results reinforce the importance of continuous adequate nutrition, especially vitamin B12, in regulation and maintenance of both nuclear DNA and mtDNA. While our data also suggests that “overnutrition”, i.e., folate excess, can also increase heteroplasmy rates, it would be premature to conclude that such overnutrition produces any adverse effects.

Uracil misincorporation has been highlighted as a potential mechanism of DNA damage in systems that have altered one-carbon metabolism. When uracil is mistakenly incorporated into DNA during replication, it can cause double-stranded breaks and ultimately cause mutations to occur^34,65^. Uracil content of the mtDNA samples were assessed in a subset of the liver samples that were also sequenced. We found higher levels of uracil in the mtDNA of all mice fed a vitamin B12 deficient diet and in *Cd320*^−/−^ mice fed the vitamin B12 replete diet. The consequences of uracil incorporation into mtDNA have not been resolved, though uracil in mtDNA is associated with impaired maximal oxidative phosphorylation capacity^66,67^. The mitochondria and the nucleus have independent DNA repair systems. It is also unclear from this research if the underlying mechanism driving uracil incorporation is shared between nuclear and mtDNA and this is an area that warrants further investigation.

The tissues of vitamin B12 deficient and excessive folic acid supplemented mice had elevated levels of mtDNA heteroplasmy compared to controls. The elevated heteroplasmy in the mice fed a vitamin B12 deficient diet replicates the findings in our first experiment, as these mice were reared, and sequencing runs were completed independently. Interestingly, folic acid supplementation appeared to cause elevated levels of mitochondrial heteroplasmy, regardless of vitamin B12 content. This effect may be larger when combined with a vitamin B12 deficient diet. Folic acid supplementation has been observed to cause elevated levels of mutagenicity in murine colon tissue, although this effect was measured in nuclear DNA^63^. Excess folic acid, at the same levels of this study, were shown to increase nuclear mutations in whole embryos. Interestingly, the authors of this study also observe the same effect in folate deficient embryos^68^. It is pertinent to note that the folic acid supplementation dosage levels used in our experiments are far above the equivalent recommended levels for humans.

Our results provide evidence that altered one-carbon metabolism can lead to increased mitochondrial heteroplasmy in mice. It is unclear, however, what mechanism(s) are driving these effects. They may include uracil misincorporation, increased ROS production or inhibition of mitochondrial DNA repair pathways. Due to the low number of variants identified across all the samples, it is not possible to assess the number of transitions, transversions or the ratio between them. A larger sample size would be required to address this question. Additional studies are needed to identify the potential mechanisms through which heteroplasmic variants occur in folic acid-supplemented and vitamin B12 deficient conditions. Similarly, the low number of heteroplasmic variants found across all samples made it unclear if there were any specific locations at which heteroplasmic variants were more likely to occur.

### Nutrient driven mtDNA heteroplasmy in a human cohort may reveal an effect of biological sex

Translating our findings from our mouse studies in a human cohort is more challenging. The FHS data were subdivided into two groups which had vitamin B12 and folate levels measured using different methodologies. There were mixed results between Group 1 and 2 when the relationship between mitochondrial heteroplasmy and folate levels was assessed. Minor allele frequency was positively correlated with folate levels across the full dataset in Group 1. This observation is in line with the results described from mice fed a folic acid supplemented chow. However, this correlation was not significant in Group 2. The lack of certainty is further confounded by the low sequencing depth in this study compared to the mouse experiments. Further studies that specifically target mitochondrial DNA sequencing are required to validate these findings as it is also possible that these results represent incidental findings, especially given the small sample size.

The number of heteroplasmic sites was not significantly affected by folate status in Group 1 but was in Group 2. This is curious, as one would expect both metrics to increase, which was observed in the mouse models. One technical explanation for this observation is that the stringency of the variant filter may have removed the ability to observe low frequency heteroplasmic alleles. This would result in a reduction of heteroplasmic sites observed in each sample, thus explaining the conflicting results between folate and number of heteroplasmic sites. Vitamin B12 concentration was found to have a significant effect on the number of heteroplasmic sites in Group 1, with lower levels of vitamin B12 correlating with a higher number of sites identified. Conversely, vitamin B12 status was not found to have a significant effect on mitochondrial heteroplasmy in Group 2.

The mixed results in human data, combined with the results observed in the mouse experiments, indicate that vitamin B12 status may also have an impact on mitochondrial DNA heteroplasmy. However, further studies are required to validate this effect. Ideally, these studies should include a specific mitochondrial DNA enrichment step in order to identify low frequency heteroplasmic sites and to accurately measure small changes in mitochondrial heteroplasmy. Additionally, one major complication of human data is the presence of genetic variation within the mitochondrial genomes of human subjects^69^. This increases the level of noise within the heteroplasmy results from the human data, which is not experienced in the mouse studies.

In conclusion, our study provides supporting evidence for the hypothesis that both folic acid and vitamin B12 status have an impact on the mutation rate of the mitochondrial genome during aging resulting in increased mitochondrial heteroplasmy. This further highlights the relevance of these micronutrients, not only during early development, but also throughout the life course.

## Materials and Methods

### Mouse Breeding and tissue harvesting

All animal protocols were reviewed and approved by the National Human Genome Research Institute (NHGRI) Animal Care and Use Committee prior to animal experiments. C57BL/6J mice were housed in shoe box cages and fed ProLab RMH 1800 diet (PMI Nutrition International) containing 50 μg vitamin B12/kg of diet and 3.3 mg folic acid/kg of diet. Breeding mice were fed Picolab Mouse Diet 20, containing 51 μg vitamin B12/kg diet and 2.9 mg folic acid/kg of diet. The diet was soy-based as vitamin B12 is not found in plant proteins, as custom formulated by Teklad Diets (Envigo, see Supplementary Table 1 for diet formulation).

#### Experimental design for mouse experiment 1

The experimental groups are displayed in Fig. 5. Briefly, *Cd320*^−/−^ and *Cd320*^+/+^ mice were divided into two dietary groups making a total of 4 experimental groups. Five mice from each of these experimental groups were used for this experiment. The mice were allowed to age on diet until reaching 1 year, at which point they were sacrificed, and tissues collected. Tissues were also harvested from a fifth group of *Cd320*^+/+^ mice maintained on a vitamin B12 replete diet after 2 years. These older tissues were used to compare the effect of vitamin B12 deficiency with the effect of aging on mitochondrial DNA heteroplasmy.

**Figure 5.**
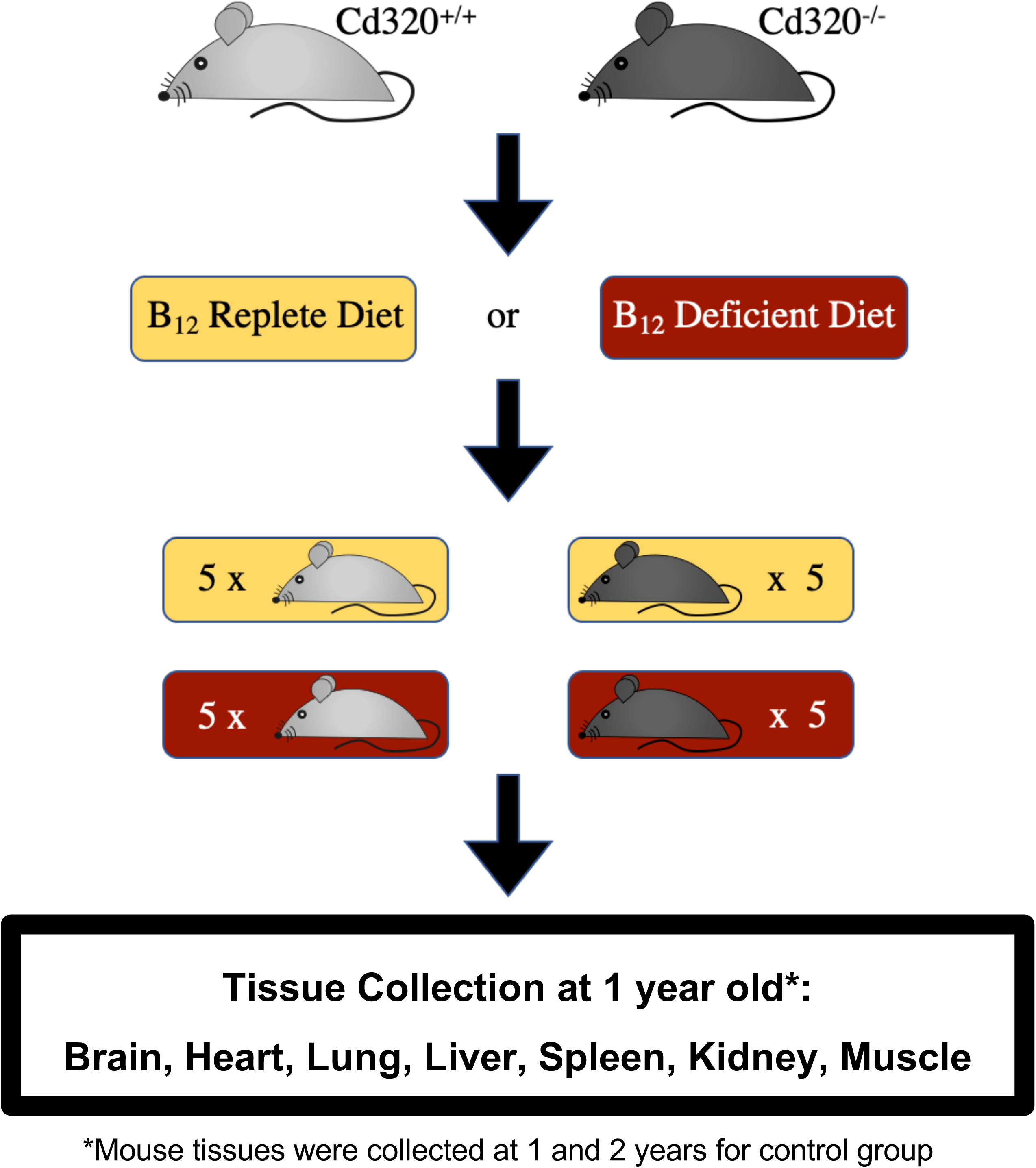
Experimental workflow for investigation of the relationship between heteroplasmy and one-carbon impaired mice. Cd320^−/−^ and Cd320^+/+^ mice were put on a cobalamin deficient or replete diet until sacrifice after 12 months. At this point the brain, heart, lungs, spleen, liver, kidney and muscle were harvested. An older cohort of Cd320^+/+^ on vitamin B12 replete diet were aged to 24 months and were also included to investigate the effect of aging on heteroplasmy. Harvested tissues underwent mitochondrial isolation and subsequent DNA extraction.

#### Experimental design for mouse experiment 2

Mice were maintained on one of four chemically-defined diets to consider the potential interactions between the status of these vitamins as previously suggested^22^. The diets included: i) vitamin B12 and folic acid ii) no vitamin B12 and folic acid iii) vitamin B12 and high folic acid and iv) no vitamin B12 and high folic acid (n=6 for each diet, Fig. 6). The make-up and concentrations of each of the diets can be found in Supplementary Table 2. Both parents were put on these diets at mating and pups were maintained on the same diet for 6 months (males, all tissues) and 12 months (males and females, liver only). At each timepoint, mice were sacrificed, and brain, heart, lung, liver, spleen, kidney and colon were collected. These samples underwent mitochondrial DNA enrichment and subsequent heteroplasmy analysis.

**Figure 6.**
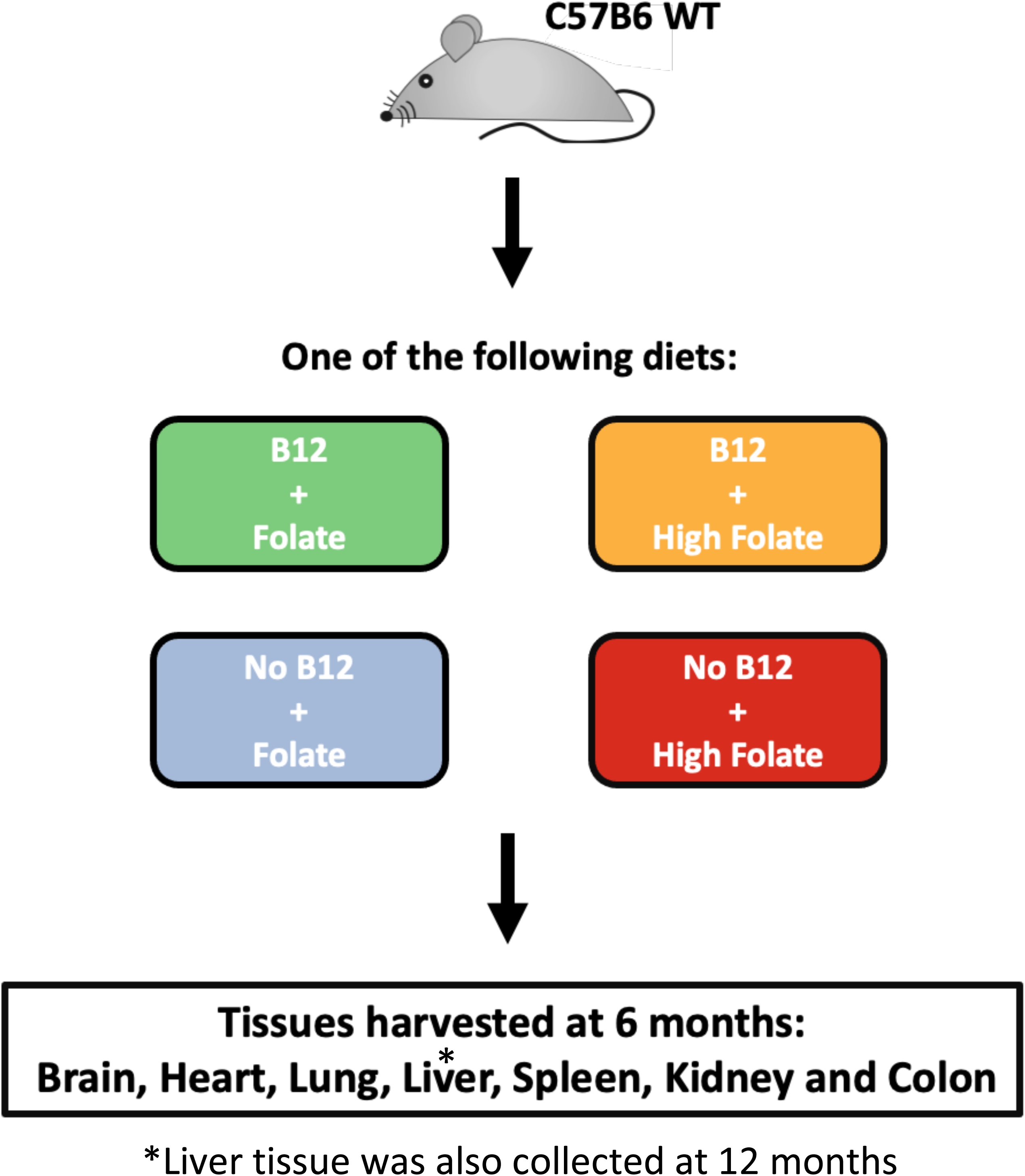
C57BL6 wild-type mice were fed one of four chemically defined diets over a 6-month period. Both parents started on the diets at mating and then pups were maintained on diet for 6 months. At this point male mice were sacrificed, and brain, heart, lung, liver, spleen, kidney and colon were isolated and underwent mitochondrial DNA isolation. Livers of both male and female mice were also harvested after 12 months.

### Tissue homogenisation

Harvested tissue was placed in a homogenization tube with 10X volume per gram of fresh homogenization buffer i.e., 5 ml buffer for 500 mg tissue. Tissues were homogenized until no discernible whole tissue was present. The homogenate was then transferred to 1.5 ml microcentrifuge tubes and spun at 1000 g for 1 minute at 4℃. The supernatant was transferred to a new microcentrifuge tube and spun at 12000 g for 10 minutes at 4℃ to pellet mitochondria. The mitochondrial pellet was resuspended with 100 µl of resuspension buffer for storage or for immediate DNA extraction.

### Mitochondrial DNA isolation

The mitochondria resuspension was added to 200 µl alkaline lysis buffer, vortexed, and placed on ice for 5 minutes. Potassium acetate buffer (150 µl) was then added, and the mixture was vortexed slowly and placed on ice for 5 minutes. The mixture was centrifuged at 12,000 g for 5 minutes at 4℃ to pellet proteins and the supernatant was decanted to a new tube. RNase (1 µg) was added to the mixture and left at room temperature for 15 minutes. Phenol-chloroform (500 µl) was added to each tube, inverted and placed on a shaker/rotator for 20 minutes. Afterwards, centrifugation at 12000 g for 2 minutes at room temperature was carried out. The aqueous (top) layer was decanted to a new tube (approx. 450 µl from this phase was retrieved) and 40 µl sodium acetate, 1 µl glycogen (20 mg/ml) and 1200 µl 100% EtOH were added. The mixture was inverted and mixed well then left on dry ice for 60 minutes. The mixture underwent centrifugation at 12,000 g and the supernatant was removed. The pellet was finally washed twice using 70% ethanol, air-dried, and resuspended in a low TE (10mM TRIS, 0.1 mMEDTA) buffer for sequencing or standard TE buffer (10mM TRIS, 1.0 mM EDTA) for (q)PCR.

### Library preparation and next generation DNA sequencing

Libraries were generated from approximately 50 ng genomic DNA using the Accel-NGS 2S Plus DNA Library Kit (Swift Biosciences) using 5 cycles of PCR to minimize PCR bias. The DNA samples were sheared by sonication (Covaris Inc., Woburn, MA) to a mean of 300 bp. Libraries were tagged with unique dual index DNA barcodes to allow pooling of libraries and minimize the impact of barcode hopping. Libraries were pooled for sequencing on the NovaSeq 6000 (Illumina) to obtain at least 7.6 million 151-base read pairs per individual library. Sequencing data was processed using RTA version 3.4.4. All sequencing data and associated metadata is available under SRA accessions PRJNA950453 and PRJNA881035.

### Data processing and alignment

As previously described^49^, FASTQ files were aligned to the mouse reference genome, GRCm38, with bwa mem using the default parameters^70^. Picard tools were used to add read groups, and to mark and remove duplicates^71^. Samtools was used to calculate the coverage across the nuclear and mitochondrial genome for each sample. Finally, R (v3.5.0) and ggplot2 (v3.3.0) were used for statistical analysis and subsequent visualization of graphs^72,73^.

The average sequencing coverage across the mitochondrial genome ranged from 40,000-120,000X and was dependent on the tissue of origin (Supplementary Fig. 5). Higher levels of coverage were observed in mtDNA from brain (n = 68), heart (n = 61), kidney (n = 59) and liver (n = 57) with lower levels in colon (n = 39), lung (n = 65), spleen (n = 67) and muscle (n = 11) (n = 329 sequenced tissue samples, Supplementary Fig. 5). The level of coverage was distributed uniformly across the mitochondrial genome except for coverage surrounding nucleotide position 0. This disruption in coverage was determined to be caused by alignment of sequencing reads to a linear reference genome as described in our previous publication^49^.

Alternative alleles across the mitochondrial genome were identified and quantified in all samples (Supplementary Fig. 6, a and b). More than 99.9% of these alleles were observed at extremely low frequencies (present in less than 0.1% of reads) and were likely caused by sequencing error. To avoid including false positive heteroplasmic calls in the alignment data, a minimum alternative allele frequency threshold of 0.2% was established (Supplementary Fig. 6, red lines). Any alternative allele that was found in less than 0.2% of reads at that locus was not deemed to be a true heteroplasmic variant for that sample. These variants were excluded from heteroplasmy analysis. The effect of raising or lowering the alternative allele frequency and base call quality thresholds on heteroplasmy analysis was also investigated (Supplementary Fig. 7). For the main sets of heteroplasmy analysis described in the results described in this paper, a phred score threshold of 30 and alternative allele frequency of 0.2% was implemented. Lowering either of these thresholds caused an increase in the number of heteroplasmic sites in all tissues. Many of these variants likely originate from sequencing errors and including them during group analysis resulted in a reduced difference in heteroplasmy observed between younger and older mouse tissues (Supplementary Fig. 7).

### mtDNA Uracil content assay

Uracil content measures were performed as described^66^. Briefly, 6 primer sets were designed to amplify the whole mitochondrial genome in 6 fragments of differing lengths. Real-time PCR was performed on each mtDNA sample in two assays: one using Phusion DNA polymerase and another using Phusion U DNA polymerase. Unlike ordinary Phusion polymerase, Phusion U does not stall at uracil in DNA and thus can copy DNA templates containing uracil. The CT shift between the two assays for each sample was used to calculate the relative levels of uracil in the DNA sample compared to a control. All measurements were compared to one control sample as a reference. The reaction conditions were as follows; pre-incubation at 95 °C for 5 min; denaturation at 95 °C for 15 s, annealing at 59 °C for 20 s and extension at 72 °C for 50 s (region 6) or 3 min (region 1-5) (40 cycles); melting at 95 °C for 5 s, 57 °C for 20 s and 95 °C continuous followed by cooling.

### Human data

IRB approval was granted for access to the sequencing data produced by the Framingham Heart Study, which was stored in the database for Genotypes and Phenotypes (dbGaP, https://www.ncbi.nlm.nih.gov/gap/). These data were produced *via* DNA sequencing performed on DNA extracted from whole blood and also from lymphoblastoid cell lines that were generated as part of the study. Lymphocytes isolated from blood were immortalized using Epstein-Barr virus to generate renewable sources of RNA and DNA. Whole exome and whole genome sequencing was performed using the Illumina HiSeq 2000 platform. The FASTQ files were downloaded to NHGRI servers from dbGaP ‘run selector’ and subsequent alignment and heteroplasmy analysis was performed.

For Group 1 individuals, non-fasting blood was drawn from each subject and frozen at −80°C. Plasma vitamin B12 concentrations were assessed using the BioRad Quantaphase II radio assay (Hercules, CA, USA;^74^. Plasma folate concentrations were also measured using the BioRad Quantaphase II radio assay (Hercules, CA, USA;^75^. These phenotypes were accessed using the dbGaP accessions phv00166968.v4.p13 and phv00166969.v4.p13.

Non-fasting blood was drawn from each subject in Group 2 upon three separate clinical visits. Plasma folate levels were measured using a microbial (*Lactobacillus casei*) assay performed in a 96-well plate and manganese supplementation^76,77^. Plasma folate measurements were also assessed using a solid-phase, no-boil radioimmunoassay in a commercial kit during visits two and three only (Diagnostic Productions Corp, Los Angeles, CA). Plasma vitamin B12 measurements were assessed once during each visit using a solid-phase, no-boil radioimmunoassay in a commercial kit (Diagnostic Productions Corp, Los Angeles, CA). These phenotypes were accessed using the dbGaP accession numbers phv0024649.v7, phv00024653.v7, phv0024663.v7, phv0024664.v7, phv0024665.v7, phv0024675.v7, phv0024676.v7 and phv0024677.v7. The average measurement was used for analysis.

## Figures and Tables

***Supplementary Figure 1. Spaghetti plots for the three heteroplasmy metrics of each tissue; number of heteroplasmic sites, average heteroplasmy and cumulative heteroplasmic burden of control mice aged 1 year old and 2 years old.*** Each line represents one individual mouse. This data presents no evidence of a tissue-specific effect on heteroplasmy in any of the three metrics investigated.

***Supplementary Figure 2. Spaghetti plots for the three heteroplasmy metrics of each mouse tissue; number of heteroplasmic sites, average heteroplasmy and cumulative heteroplasmic burden for the experimental groups.*** Each line represents one individual mouse. This data presents no evidence of a tissue-specific effect on heteroplasmy in any of the three metrics investigated.

***Supplementary Figure 3. Higher levels of heteroplasmy found in mice on altered diets did not show a tissue-specific pattern.*** Mice fed vitamin B12 deficient, or folic acid-supplemented diets had elevated variant frequencies across all tissues and did not show a tissue-specific pattern (n=6 for each tissue and group).

***Supplementary Figure 4. Mitochondrial heteroplasmy was also elevated in livers of male and female mice maintained on vitamin B12 deficient and folic acid-supplemented diets for 12 months.*** There was a significant relationship between the number of heteroplasmic variants/average heteroplasmy and diet (Kruskal-Wallis). More variants were found in the tissues of mice that were fed a vitamin B12 deficient and/or folic acid-supplemented diet. The y axis represents the a, number of heteroplasmic sites, b, average heteroplasmy and c, cumulative heteroplasmic burden. The diets are plotted on the x axis.

***Supplementary Figure 5. Read coverage across the mitochondrial genome after sequencing and subsequent data-processing.*** The average sequencing depth ranged from 40,000-120,000X. Higher levels of coverage were observed in mtDNA from brain (n = 68), heart (n = 61), kidney (n = 59) and liver (n = 57) with lower levels in colon (n = 39), lung (n = 65), spleen (n = 67) and muscle (n = 11) (n = 329 sequenced tissue samples). Brain, heart, kidney and liver had similar levels of sequencing coverage across the genome, with lower levels found in lung, muscle and spleen. A fluctuation in coverage was observed at nucleotide position 0 when the default reference genome was used and was due to alignment to a linear reference in contrast to the circular nature of the mitochondrial genome.

***Supplementary Figure 6. Alternative alleles and their frequency observed across the mitochondrial genome in all sequenced samples.*** The vast majority of alternative alleles observed were likely due to sequencing error and thus a threshold of 0.2% (red line) was implemented to exclude any variants that were likely caused by sequencing error. The green line signifies an alternative allele frequency of 10%. There were only 5 variants found above this frequency across all samples in the first mouse experiment and 12 were found in the second.

***Supplementary Figure 7. Larger differences in heteroplasmic sites and average alternative allele frequency are observed between young and old mice when quality and alternative allele frequency thresholds are implemented.*** Phred quality score threshold increases from left to right. Alternative allele frequency threshold increases from top to bottom. When lower quality base-calls are included in heteroplasmy analysis, more false positive alternative allele calls are introduced into analysis. This is also the case for including heteroplasmic variants at lower frequencies, which leads to smaller differences in heteroplasmy between younger and older mouse tissues.

**Supplementary Table 1 and 2.**
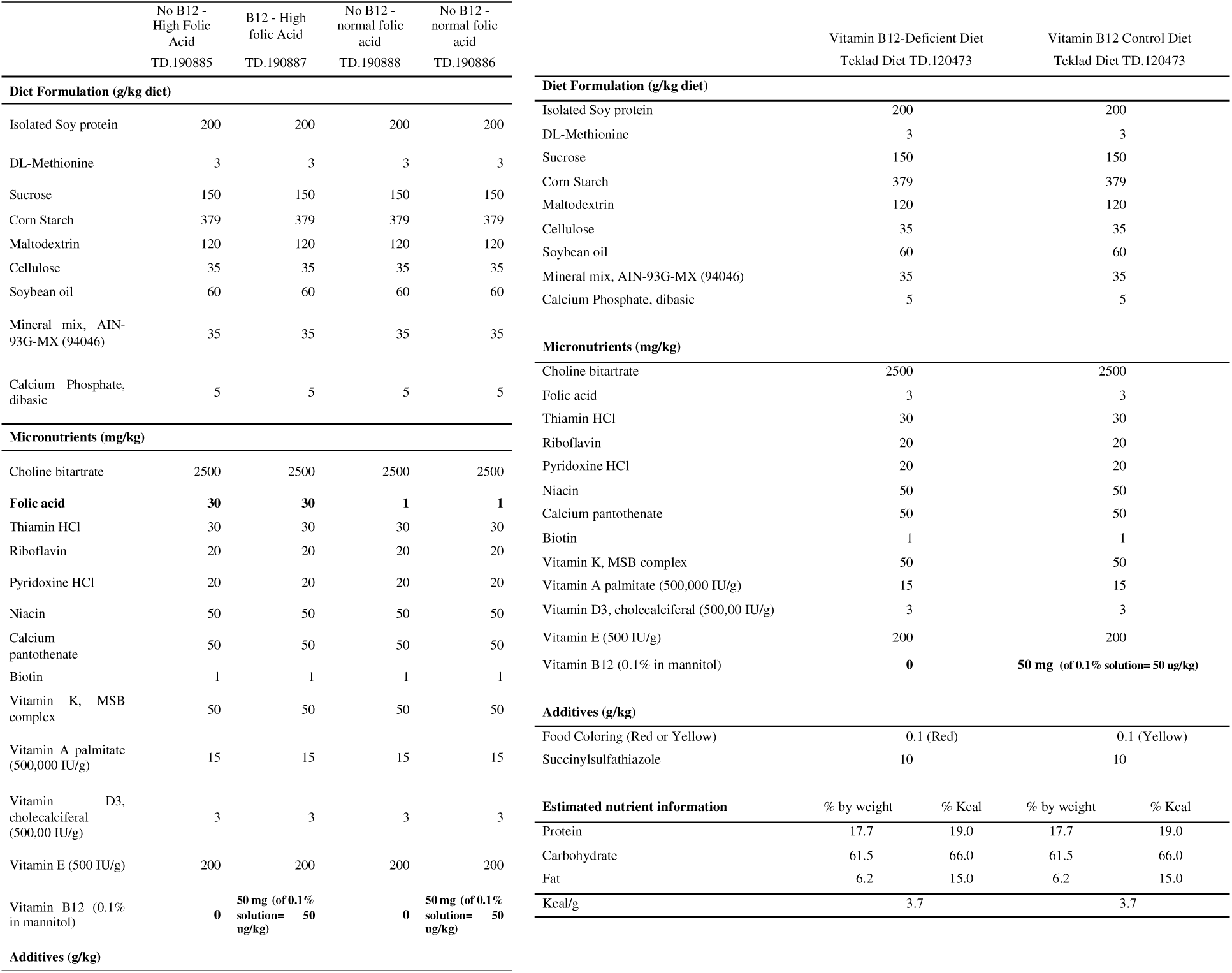
Custom diet formulations used in both experiments. Note that the vitamin B12 is in a 0.1% solution, such that the inclusion of 50 mg of the 0.1% solution equals 50 µg vitamin B12/kg of diet.

**Supplementary Table 3.** The number and breakdown by sex of individuals in Groups 1 and 2 with biochemical measurements and sequencing data.

|  | *Male (n)* | *Female (n)* | *Total (n)* |
| --- | --- | --- | --- |
| **Group 1** |  |  |  |
| *Blood measurements* |  |  | 1991 |
| *With WGS* | 16 | 18 | 34 |
| *With WXS* | 15 | 9 | 24 |
| *Total* | 31 | 27 | 58 |
| **Group 2** |  |  |  |
| *Blood measurements* |  |  | 2612 |
| *With WGS* | 150 | 143 | 293 |
| *Total* | 150 | 143 | 293 |
| *Sample type* |  |  |  |
| *Blood* | 85 | 58 | 143 |
| *Cell line* | 75 | 75 | 150 |

## Notes

https://www.ncbi.nlm.nih.gov/bioproject

### Competing Interest Statement

The authors have declared no competing interest.

