## Supplementary material for "Vitamin B12 status and folic acid supplementation influence mitochondrial heteroplasmy levels in mice as they age": Suppl

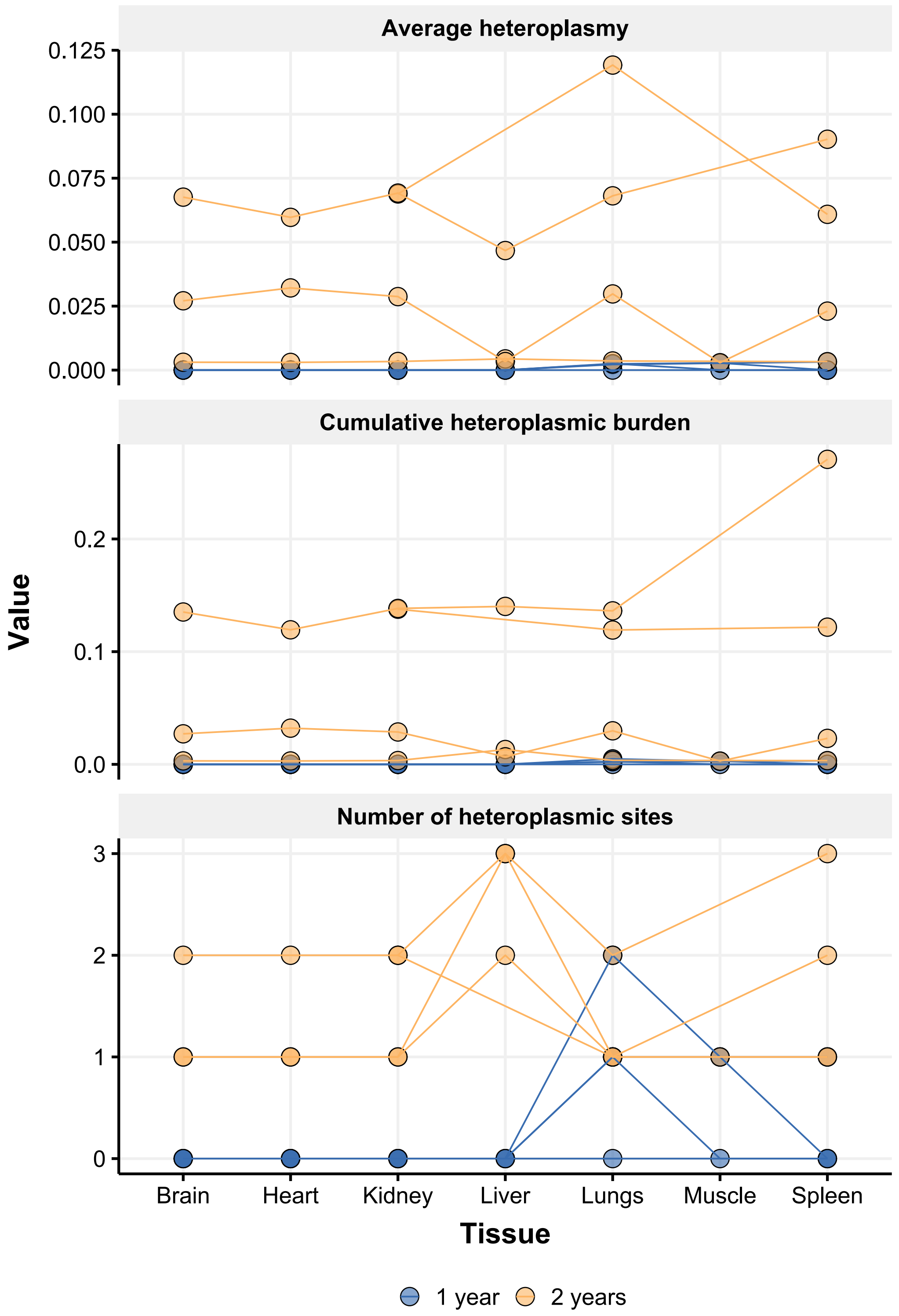


***Supplementary Figure 1. Spaghetti plots for the three heteroplasmy metrics of each tissue; number of heteroplasmic sites, average heteroplasmy and cumulative heteroplasmic burden of control mice aged 1 year old and 2 years old.*** Each line represents one individual mouse. This data presents no evidence of a tissue-specific effect on heteroplasmy in any of the three metrics investigated.

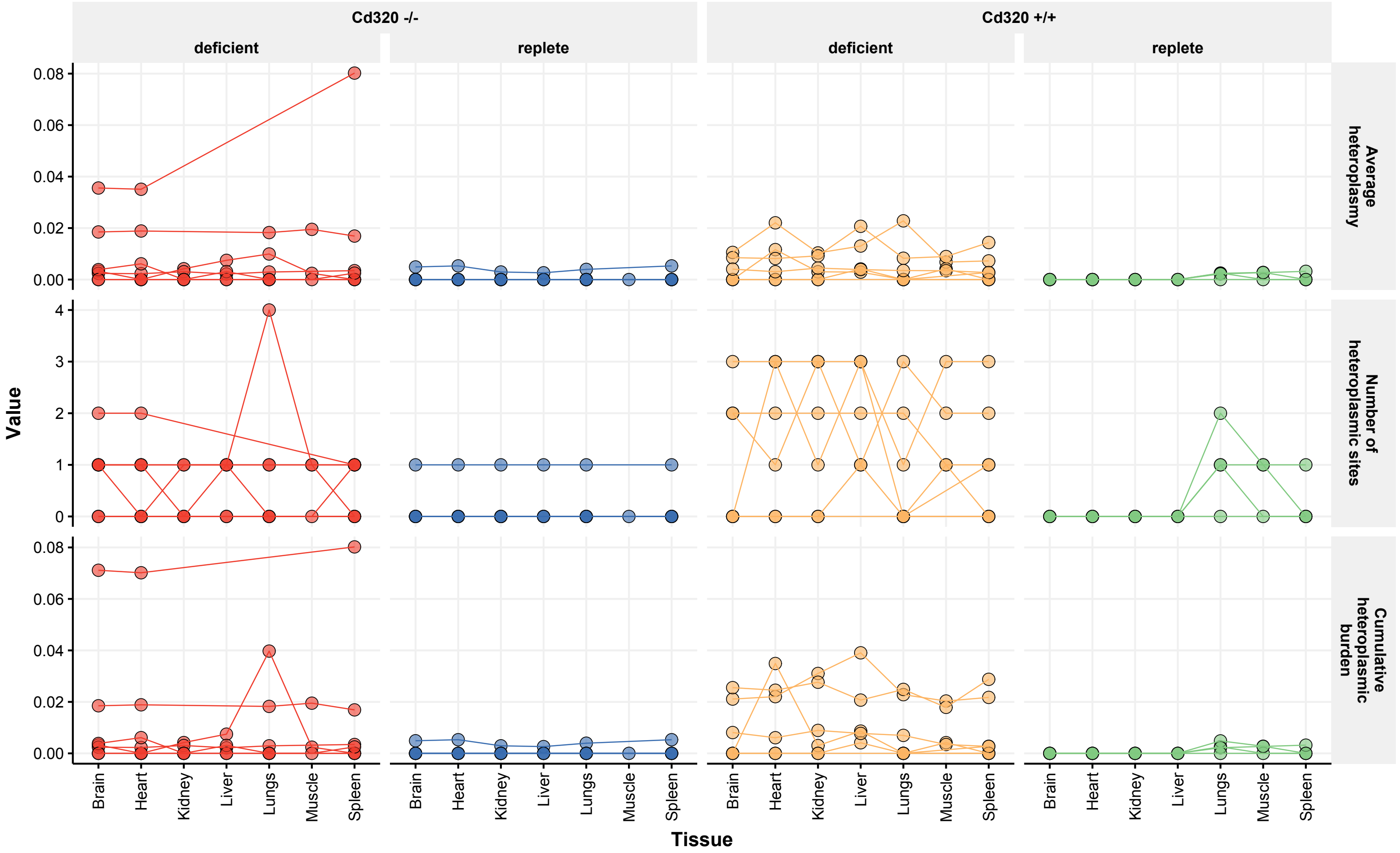


***Supplementary Figure 2. Spaghetti plots for the three heteroplasmy metrics of each mouse tissue; number of heteroplasmic sites, average heteroplasmy and cumulative heteroplasmic burden for the experimental groups.*** Each line represents one individual mouse. This data presents no evidence of a tissue-specific effect on heteroplasmy in any of the three metrics investigated.

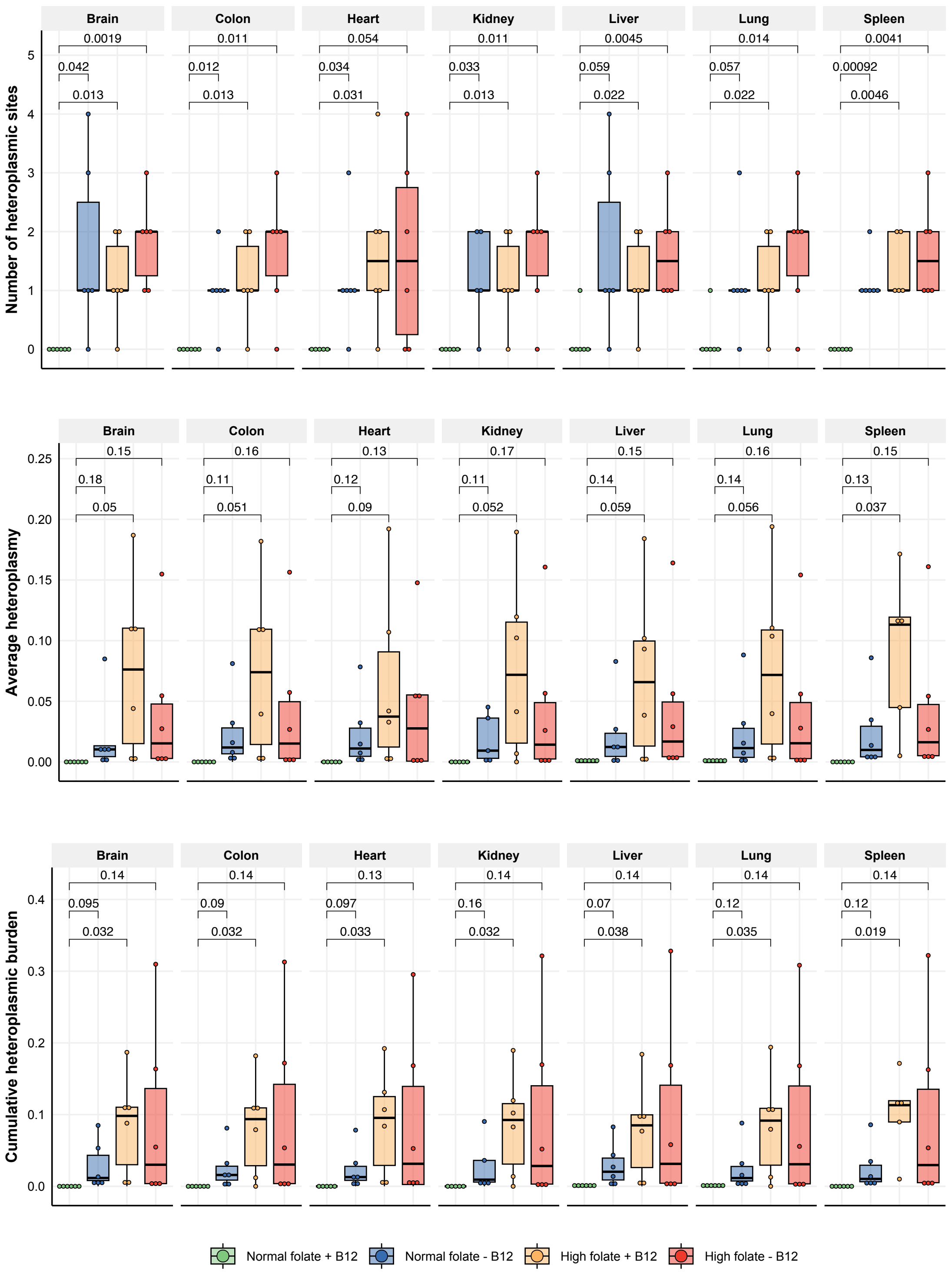


***Supplementary Figure 3. Higher levels of heteroplasmy found in mice on altered diets did not show a tissue-specific pattern.*** Mice fed vitamin B12 deficient, or folic acid-supplemented diets had elevated variant frequencies across all tissues and did not show a tissue-specific pattern (n=6 for each tissue and group).

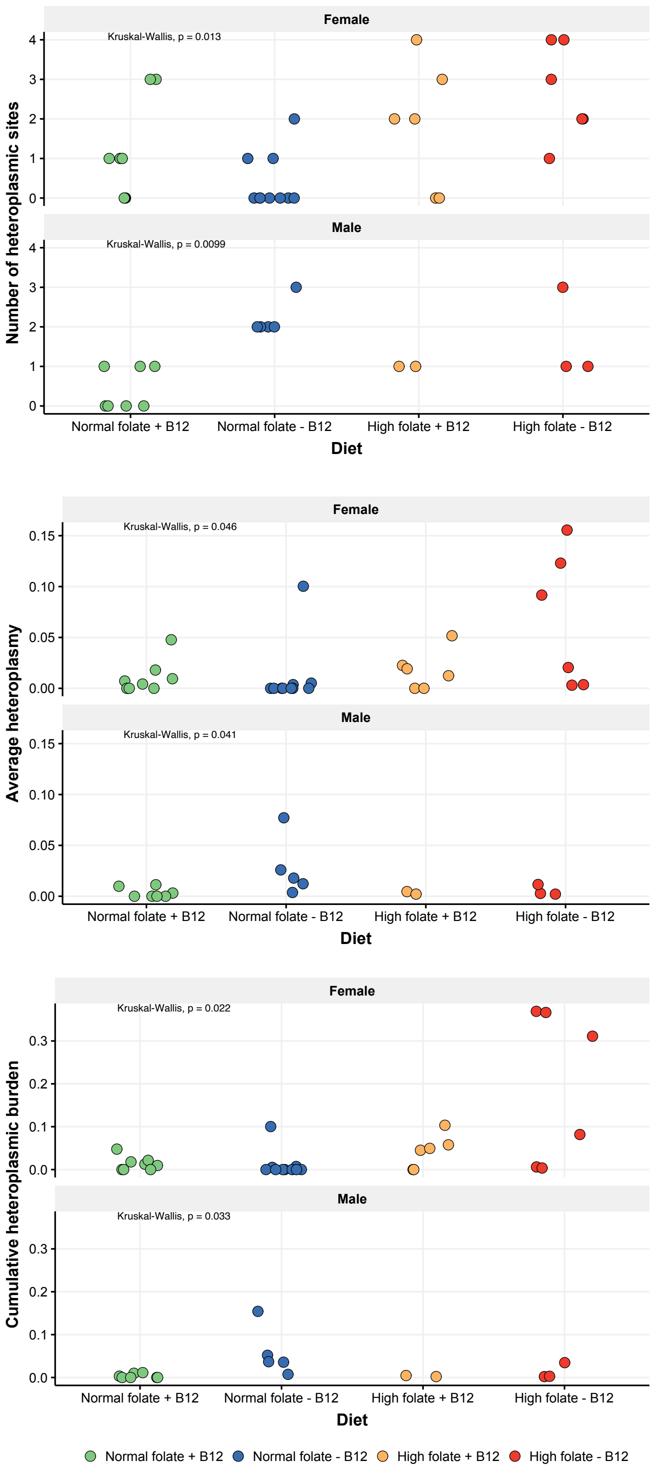


***Supplementary Figure 4. Mitochondrial heteroplasmy was also elevated in livers of male and female mice maintained on vitamin B12 deficient and folic acid-supplemented diets for 12 months.*** There was a significant relationship between the number of heteroplasmic variants/average heteroplasmy and diet (Kruskal-Wallis). More variants were found in the tissues of mice that were fed a vitamin B12 deficient and/or folic acid-supplemented diet. The y axis represents the a, number of heteroplasmic sites, b, average heteroplasmy and c, cumulative heteroplasmic burden. The diets are plotted on the x axis.

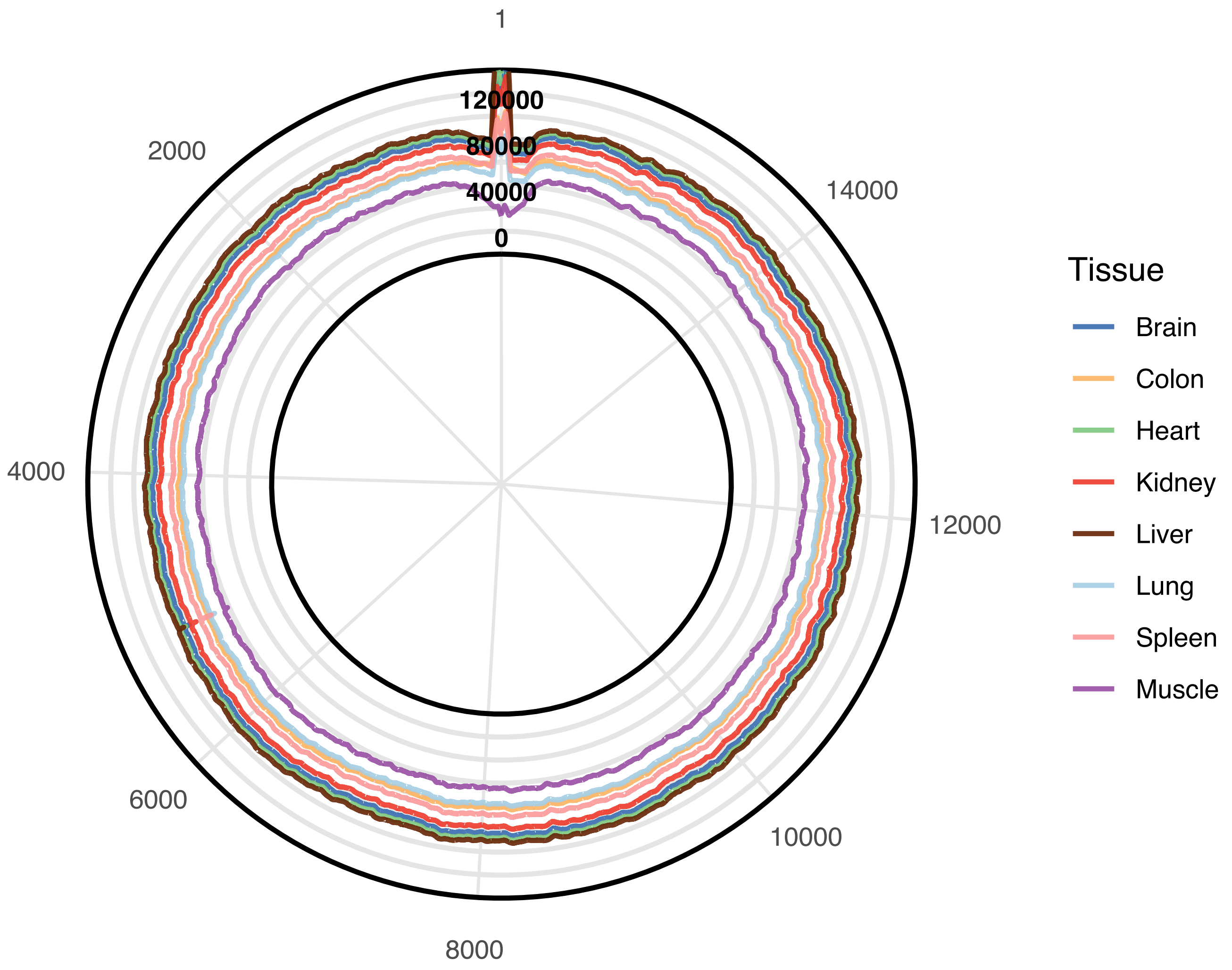


***Supplementary Figure 5. Read coverage across the mitochondrial genome after sequencing and subsequent data-processing.*** The average sequencing depth ranged from 40,000-120,000X. Higher levels of coverage were observed in mtDNA from brain (n = 68), heart (n = 61), kidney (n = 59) and liver (n = 57) with lower levels in colon (n = 39), lung (n = 65), spleen (n = 67) and muscle (n = 11) (n = 329 sequenced tissue samples). Brain, heart, kidney and liver had similar levels of sequencing coverage across the genome, with lower levels found in lung, muscle and spleen. A fluctuation in coverage was observed at nucleotide position 0 when the default reference genome was used and was due to alignment to a linear reference in contrast to the circular nature of the mitochondrial genome.

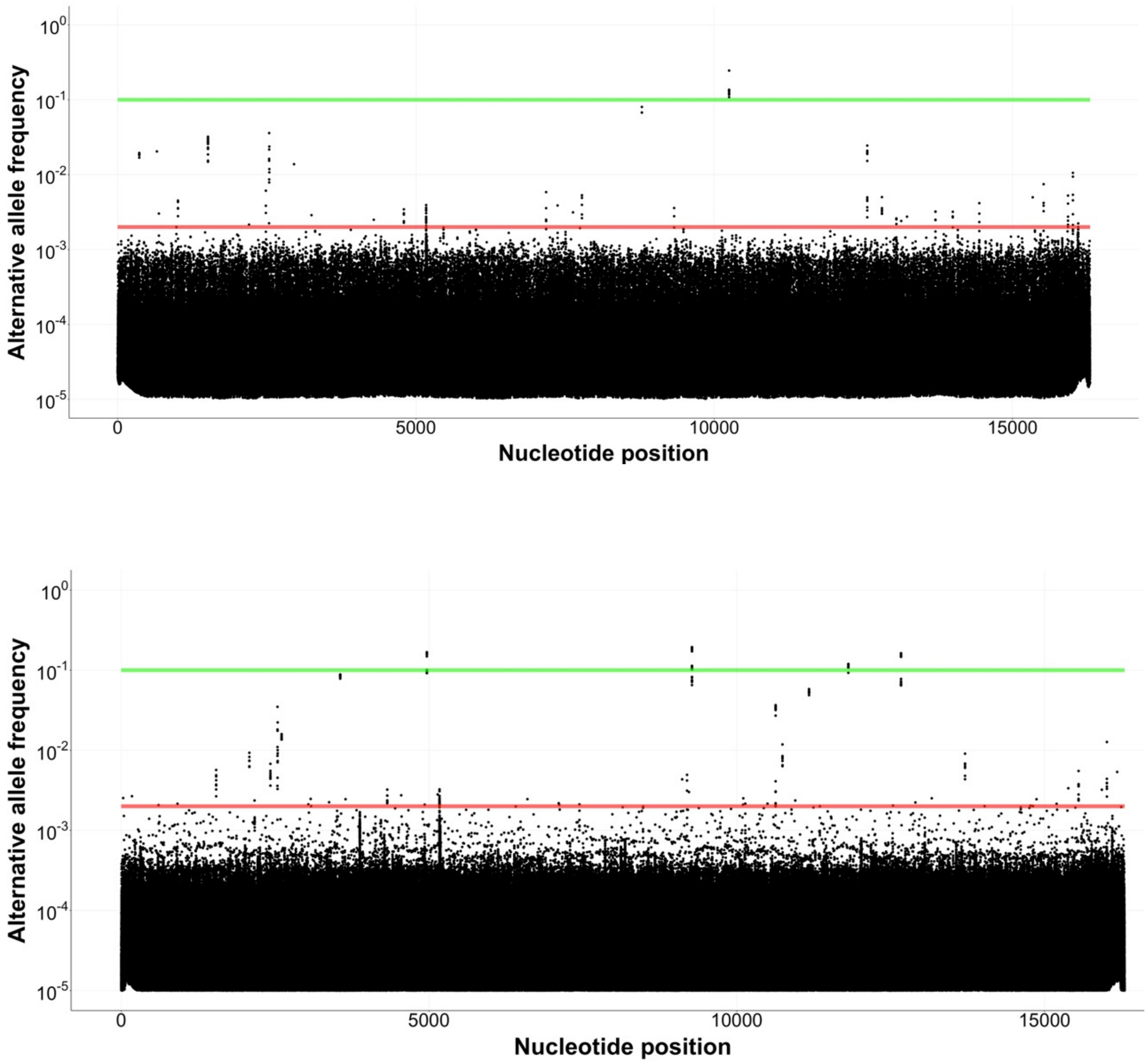


***Supplementary Figure 6. Alternative alleles and their frequency observed across the mitochondrial genome in all sequenced samples.*** The vast majority of alternative alleles observed were likely due to sequencing error and thus a threshold of 0.2% (red line) was implemented to exclude any variants that were likely caused by sequencing error. The green line signifies an alternative allele frequency of 10%. There were only 5 variants found above this frequency across all samples in the first mouse experiment (top) and 12 were found in the second (bottom).

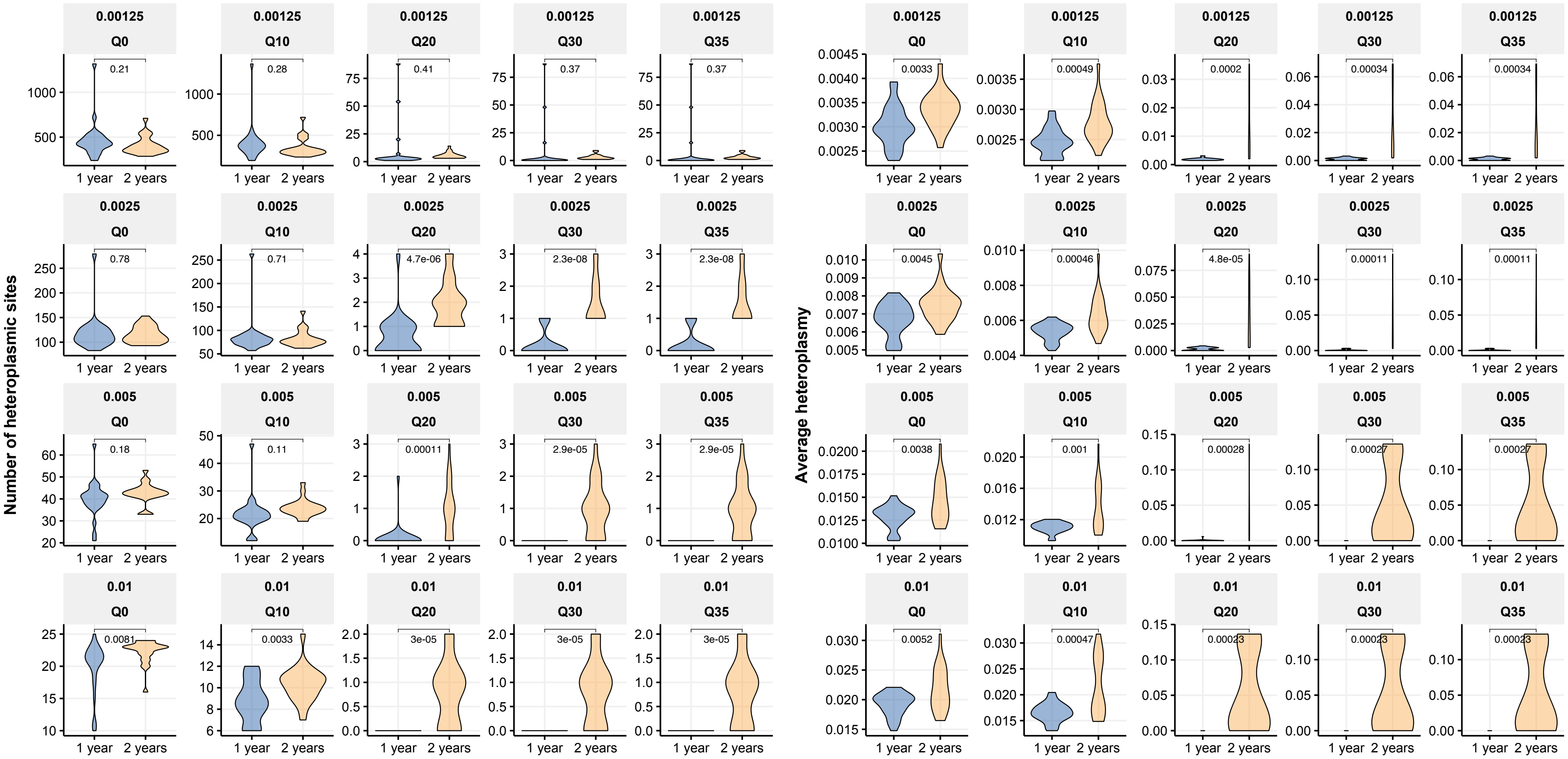


***Supplementary Figure 7. Larger differences in heteroplasmic sites and average alternative allele frequency are observed between young and old mice when quality and alternative allele frequency thresholds are implemented.*** Phred quality score threshold increases from left to right. Alternative allele frequency threshold increases from top to bottom. When lower quality base-calls are included in heteroplasmy analysis, more false positive alternative allele calls are introduced into analysis. This is also the case for including heteroplasmic variants at lower frequencies, which leads to smaller differences in heteroplasmy between younger and older mouse tissues.
